# A Global Transcriptional Atlas of the Effect of Sleep Deprivation in the Mouse Frontal Cortex

**DOI:** 10.1101/2023.11.28.569011

**Authors:** Kaitlyn Ford, Elena Zuin, Dario Righelli, Elizabeth Medina, Hannah Schoch, Kristan Singletary, Christine Muheim, Marcos G. Frank, Stephanie C. Hicks, Davide Risso, Lucia Peixoto

**Author notes:** Authors contributed equally (Kaitlyn Ford, Elena Zuin).

## Abstract

Sleep deprivation (SD) has negative effects on brain function. Sleep problems are prevalent in neurodevelopmental, neurodegenerative and psychiatric disorders. Thus, understanding the molecular consequences of SD is of fundamental importance in neuroscience. In this study, we present the first simultaneous bulk and single-nuclear (sn)RNA sequencing characterization of the effects of SD in the mouse frontal cortex. We show that SD predominantly affects glutamatergic neurons, specifically in layers 4 and 5, and produces isoform switching of thousands of transcripts. At both the global and cell-type specific level, SD has a large repressive effect on transcription, down-regulating thousands of genes and transcripts; underscoring the importance of accounting for the effects of sleep loss in transcriptome studies of brain function. As a resource we provide extensive characterizations of cell types, genes, transcripts and pathways affected by SD; as well as tutorials for data analysis.

## INTRODUCTION

Sleep is an evolutionary conserved powerful drive, but its function remains a mystery. It is well established that sleep deprivation (SD) has negative effects on brain function and affects a wide array of molecular processes^1^. Sleep problems are widely observed in neurodevelopmental, neurodegenerative and psychiatric disorders^2,3^. Thus, understanding the molecular consequences of SD is of fundamental importance in neuroscience. We and others have shown that in rodents SD strongly affects the brain transcriptome^4–13^. It was initially thought that there was little agreement on the effects of SD. However, we have shown that if biological and technical noise are properly accounted for, hundreds of genes are differentially expressed after SD in the mouse brain regardless of technology, site or brain region^14^. Currently, studies of the effect of SD on the brain transcriptome are focused on genes. However, in mammals cells often express multiple transcripts of the same gene (isoforms). Also, most current studies also lack resolution at the cell type level with sufficient statistical power.

To further our understanding of the molecular consequences of acute SD, in this study we present the first simultaneous bulk and single-nuclear (sn)RNA sequencing characterization of the effects of SD in the frontal cortex of adult male mice at high resolution. We chose the frontal cortex because in humans it is the brain region most strongly affected by SD^15^. Using snRNA-seq we show that SD predominantly affects glutamatergic neurons, specifically in layers 4 and 5. Using bulk RNA-seq we performed differential gene and transcript expression (DGE/DTE), as well as differential transcript usage (DTU) analysis. We show that at the bulk level SD affects half of the frontal cortex transcriptome and produces isoform switching of thousands of transcripts. Both bulk and snRNA-seq analysis show that SD has a large repressive effect on transcription, down-regulating thousands of genes and transcripts both globally and in specific cell types. This large yet cell-specific effect underscores the importance of controlling or accounting for the effects of sleep loss in any transcriptome studies of brain function. As a resource to the neuroscience community we provide extensive characterization of which genes, transcripts and pathways are affected by SD and in which cell types; as well as guided tutorials for reproducible snRNA-seq differential expression, ask well as bulk-RNA-seq simultaneous DGE, DTE and DTU analysis.

## RESULTS

### Sleep deprivation preferentially affects gene expression in neurons

It is currently unknown how different cell types respond to sleep deprivation (SD) in the mouse frontal cortex, and which genes and pathways are differentially affected across different types of neurons (e.g. glutamatergic and GABAergic). To address this, we carried out single-nuclear (sn)RNA-seq using 10X Genomics Chromium v3 technology followed by Illumina sequencing of adult male mice either allowed to sleep in their home cages (HC) or sleep deprived (SD, n=3 per group). After gene abundance quantification, we performed cell type label assignment using a reference dataset obtained from the Brain Initiative cell Census Network (BICCN) to avoid the lack of reproducibility of cell-type labels that can arise from cluster-based assignment ^16^. Nuclei counts for each replicate for each cell type are available in Extended Data Figure 1. Cell type nomenclature follows BICCN guidelines as per Yao et al., 2021. Glutamatergic neurons are labeled based on the layer in which they reside (L1-6) as well as where they project. Across the cortex, there are three main classes of projection neurons: IT (intratelencephalic tract), PT (pyramidal tract) and CT (Corticothalamic tract). GABAergic neurons are labeled based on marker expression (e.g Parvalvumin, Somatostatin, ViP). Figure 1A depicts a UMAP visualization of the top principal components after transfer of all cell type labels in all biological replicates (3 HC, 3 SD). We then tested the robustness of cell type assignments. First, we used an independent dataset from the BICCN as a gold standard and show that our pipeline shows good consistency with known cell labels when the Yao et al., 2021 dataset is used as a reference (Extended Data Figure 2) even when two different methods are used (Azimuth and SingleR). This indicates that the reference dataset seems to be the most important aspect when assigning cell-type labels. Subsequently, to further validate the cell-type assignments in our datasets, we created heatmaps using known mouse cortex cell-specific marker expression in each cell type, showing good consistency between marker expression and cell-type assignment (Extended Data Figure 3).

**Figure 1.**
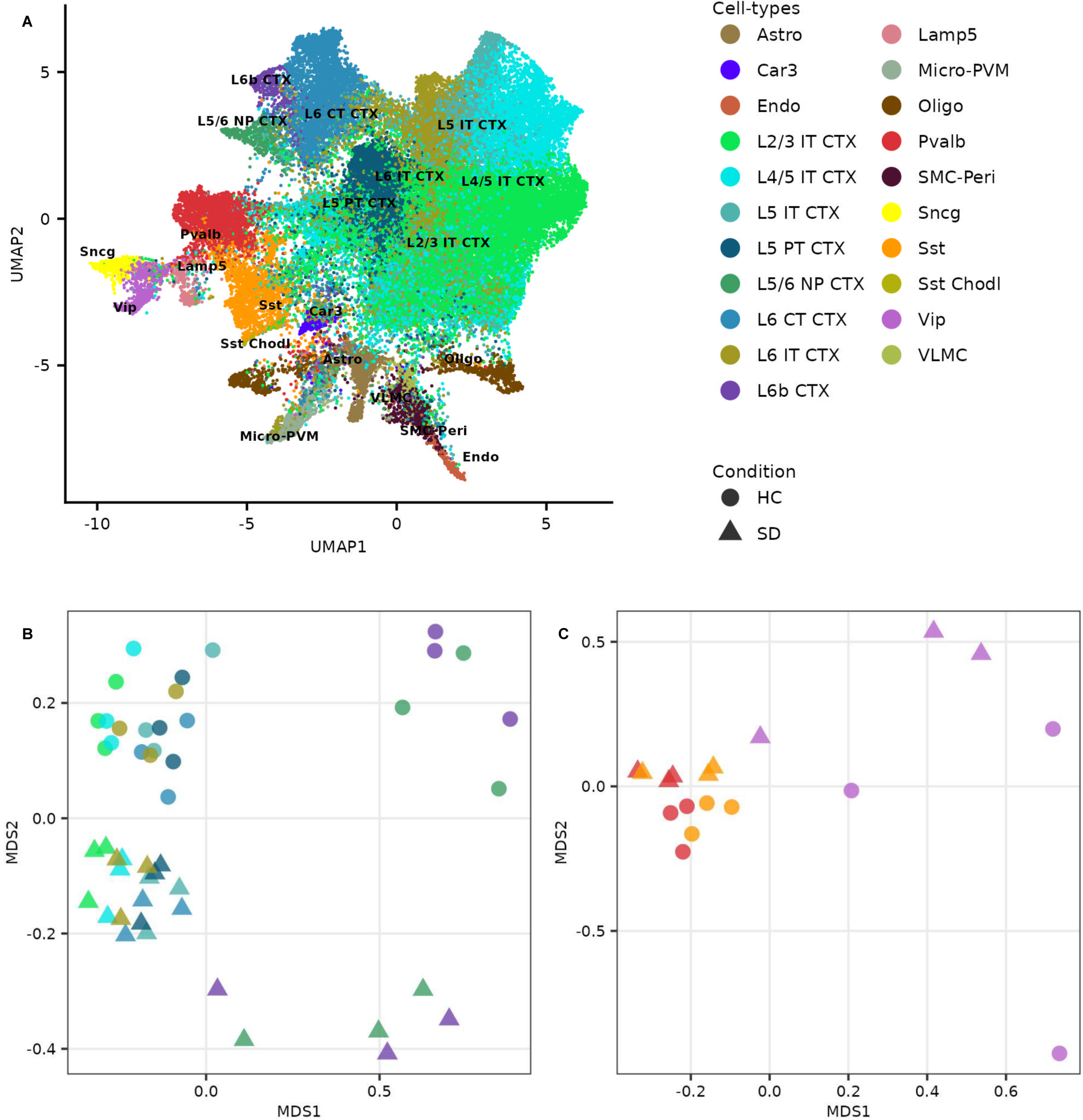
SD has a large detectable effect on glutamatergic and GABAergic cell-types. **A)** Uniform Manifold Approximation and Projection (UMAP) plot for the 52,651 frontal cortex nuclei from all 6 replicates, annotated with a reference-based Azimuth method^16^, Brain Initiative Cell Census Network, BICCN). Glutamatergic, GABAergic and non-neuronal cell-types are shaded in unique colors. **B)** Multidimensional scaling (MDS) plot for glutamatergic cell-types with more than 500 nuclei shows cell-types are separated along MDS1, accounting for the largest source of dissimilarity in the data, while HC (circles, N=3) and SD (triangles, N=3) are separated along MDS2, accounting for the remainder of dissimilarity in the data. **C)** MDS plot for GABAergic cell-types with more than 500 nuclei shows cell-types are separated along MDS1, accounting for the largest source of dissimilarity in the data, while HC (circles, N=3) and SD (triangles, N=3) are separated along MDS2, accounting for the remainder of dissimilarity in the data. HC, Home Cage Controls. SD, Sleep Deprived.

To uncover which factors drive differences in gene expression after SD across cell types, we performed multidimensional scaling (MDS) of the pseudo-bulk sum of all nuclei assigned to a cell type per independent biological replicate (color-coded by cell type and shaped by condition, HC as circles, SD as triangles). Our data shows that SD has a larger effect on glutamatergic (Figure 1B) than GABAergic neurons (Figure 1C). SD does not seem to cause major differences in gene expression in non-neuronal cell types such as Astrocytes, Oligodendrocytes or Microglia (Extended Data Figure 4A). Differences between conditions cannot be detected if we restrict the MDS plot to genes less likely to be differentially expressed by SD according to publicly available data (Extended Data Figure 4B). A tutorial for mouse brain reference-based cell-type assignment is available through GitHub and at the following website: https://rissolab.github.io/AtlasCortexSD/

### Sleep deprivation disproportionally affects glutamatergic neurons of the deeper layers of the cortex, particularly layer 4/5

To define differentially expressed genes (DEGs) after SD in each neuronal cell-type we performed pseudo-bulk analysis after normalization to remove unwanted variation^17,18^. Figure 2A summarizes, for each cell type, the number of nuclei, DEGs and positive controls, defined on the basis of publicly available data (see Online Methods). A full list of DEGs per cell type is available in Figure 2 Supplementary Table 1. To determine which cell types were disproportionately affected by SD, we compared the proportion of DEGs relative to the abundance (number of nuclei) of each cell-type (Figure 2B). Since the number of DEGs is expected to increase linearly with the number of nuclei sequenced due to the distributional properties of count models, cell-types above the line are affected more than expected by SD, while those below the line are affected less than expected. Consistent with our MDS plots, the number of DEGs was only more than expected in glutamatergic cell-types, specifically in neurons in layers 4/5 and 5 that project to the intra-telencephalic tract (L4/5 IT and L5 IT) and neurons in layers 6 (L6b).

**Figure 2.**
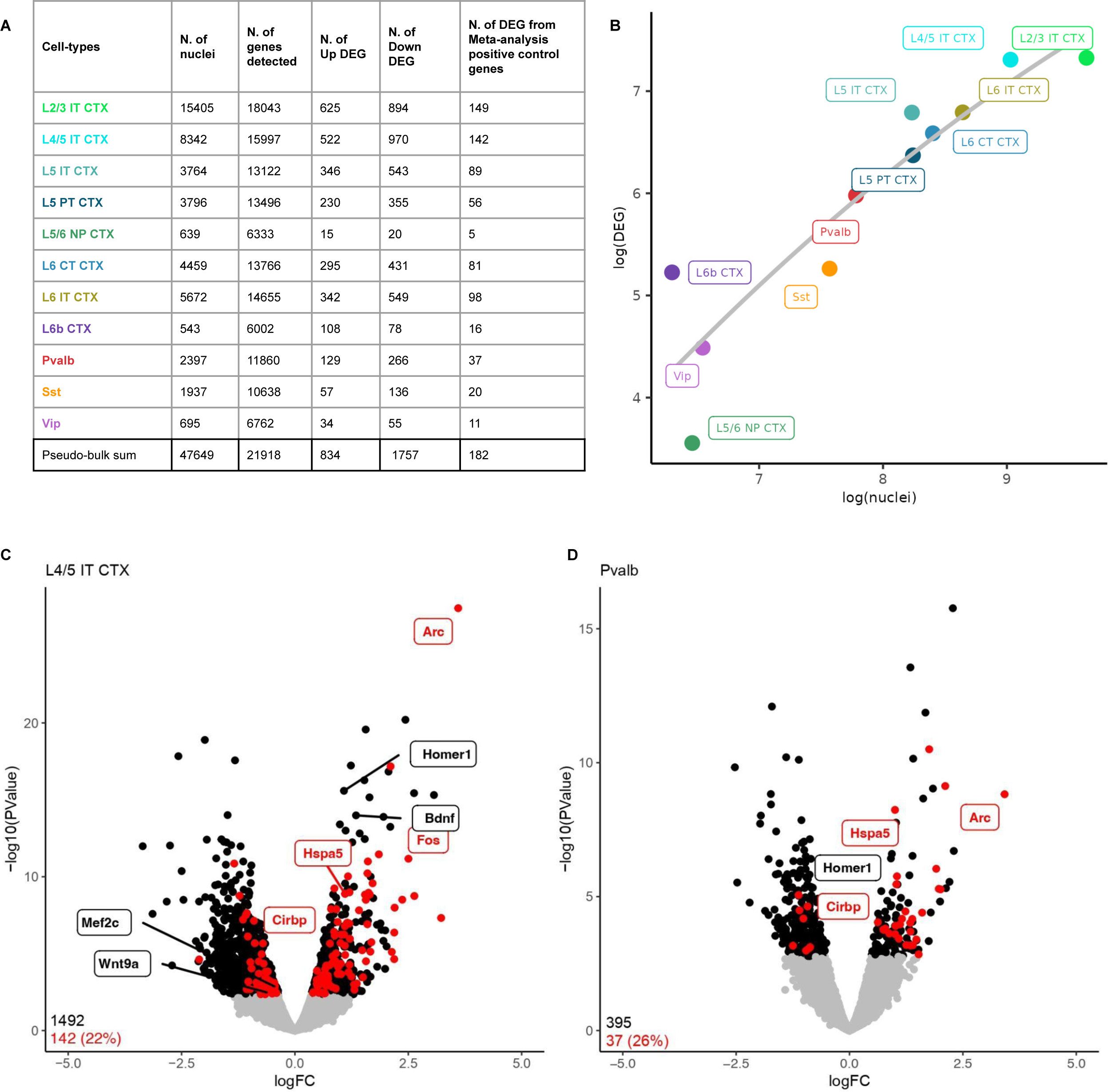
SD predominantly affects deep layers of glutamatergic cell-types. **A)** Table shows the total number of nuclei in a neuronal cell-type, the number of genes detected, and up-and down-regulated differentially expressed genes (DEGs). Also shown are the recovery of positive control genes from a prior, independent analysis^14^. The bottom row shows the pseudo-bulk sum for the expression analysis. **B)** Scatter plot with the log of the total number of nuclei on the x-axis and the log of the DEGs on the y-axis. A line of best fit was drawn through the points, with cell-types that appear to be more affected by the treatment above the line, and cell-types that appear to be less affected by the treatment below the line. **C)** Volcano plot for L4/5 IT CTX shows the logFC on the x-axis and the -log10 of the p-value on the y-axis. Expressed genes (13,592) are in gray. Significantly differentially expressed genes are in black (2,405), FDR < 0.05 and positive controls are shown in red (177). **D)** Volcano plot for Pvalb shows the logFC on the x-axis and the -log10 of the p-value on the y-axis. Expressed genes (11,585) are in gray. Significantly differentially expressed genes are in black (275), FDR < 0.05 and positive controls are shown in red (32). A subset of genes from Figure 2 Supplementary Table 2 are shown in **C** and **D**. N=3 per condition. SD, Sleep Deprivation.

Glutamatergic neurons of L4/5 IT, which express high levels of Ror-beta (Extended Data Figure 3) were the most affected with 1492 DEGs (522 upregulated, 970 downregulated, and 142 positive controls, Figure 2C). In contrast, the most affected of the GABAergic neurons, Pvalb interneurons only had 395 DEGs after SD (129 upregulated, 266 downregulated, 37 positive controls, Figure 2D). Most cell types had more down regulated than upregulated DEGs, thus SD seems to disproportionately repress transcription. Principal component analysis (PCA) and histograms of uncorrected *p*-values to ensure accurate normalization as well as volcano plots of DEG per cell type are available in Extended Data Figures 5 and 6. A tutorial for pseudo-bulk differential gene expression analysis in response to treatment of sc/snRNA-seq data is available through GitHub and at the following website: https://rissolab.github.io/AtlasCortexSD.

### Sleep deprivation affects distinct pathways and molecular functions in glutamatergic and GABAergic neurons

To better understand which genes and pathways are shared between or unique across cell-types we first intersected the union of all DEGs in glutamatergic neurons with the union of all DEGs in GABAergic neurons (Figure 3, Figure 3 Supplementary Table 1). Glutamatergic neurons contain almost 20-fold more DEGs after SD that are unique relative to GABAergic neurons (2236 vs. 109). The majority of DEGs in GABAergic neurons are shared with glutamatergic neurons (416) and contain immediate early genes (IEGs, *Arc*, *Homer1*, *Bdnf*) and stress response genes (*Hspa5*, *Hspa8*, Figure 3A). To further understand which pathways (KEGG), molecular functions (MP, Interpro) and biological processes (BP, Interpro) were affected by SD in GABAergic versus glutamatergic neurons more often than expected by chance, we carried out functional enrichment analysis (Figure 3B and 3C, Figure 3 Supplementary Table 2) of the unique sets of DEGs. When multiple molecular functions (MF), biological processes (BP) or pathways (KEGG) had overlapping sets of genes, they were clustered for ease of interpretation (see methods). Neurotransmitter receptors were downregulated in response to SD in both glutamatergic and GABAergic neurons (KEGG pathway: Neuroactive-ligand receptor interaction, *Vip*, *Sst*, *Gria4*, *Grin2d*). However, neurogenesis (Figure 3C Cluster 1 in red, BP and MF), Cell adhesion (Figure 3C Cluster 2 in red, BP and KEGG), MAPK-Akt-PI3K signaling (Figure 3C Cluster 3 in red, *MF and KEGG*), circadian rhythms (KEGG in red) were enriched in genes upregulated by SD. Genes that belong to development and differentiation (Figure 3C Cluster 1 in blue, BP and MF) were downregulated by SD only in glutamatergic neurons.

**Figure 3.**
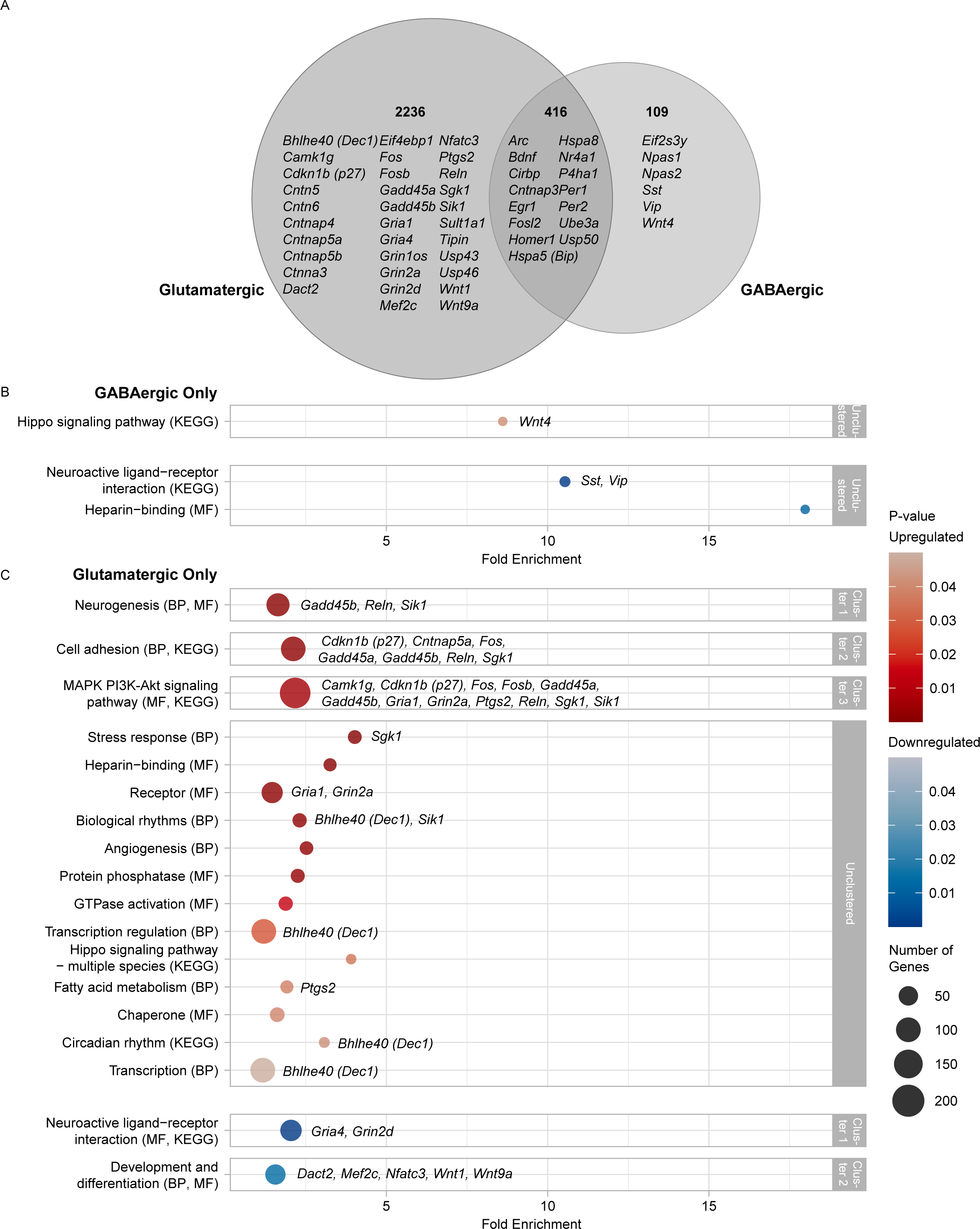
Glutamatergic neurons are preferentially affected by SD. **A)** Venn diagram of differentially expressed genes (DEGs) within glutamatergic and GABAergic neuronal cell-types (FDR < 0.05, > 500 nuclei). Genes from Figure 2 Supplementary Table 2 are highlighted. **B-C)** Bubbles show enriched terms (modified Fisher’s Exact p-value < 0.05) of upregulated (red) and downregulated (blue) DEGs that are unique to **B)** GABAergic or **C)** glutamatergic neurons, as compared to the union of expressed genes within the respective category. Bubble size reflects the number of genes per term (minimum of 3). Gray boxes outline clustered terms (similarity threshold > 0.2, and enrichment score > 1.5). Enrichment scores for each cluster; Cluster 1 Up (2.99), Cluster 2 Up (2.69), Cluster 3 Up (2.23), Cluster 1 Down (3.66), Cluster 2 Down (1.87). Terms were intersected with genes from Figure 2 Supplementary Table 2. N=3 per condition. SD, Sleep Deprivation. BP, Uniprot biological process. MF, Uniprot molecular function. KEGG, Kegg pathways.

We then asked which pathways (KEGG), molecular functions (MP) and biological processes (BP) were affected by SD more often than expected by chance only in some cell types (Figure 4, Figure 4 Supplementary Table 1). We first determined which DEGs were unique to each cell-type (Figure 4A). Glutamatergic neurons had the largest numbers of unique DEGs, specifically those in layers 2/3 and 4/5 that project to the intratelencephalic tract (L4/5 IT and L2/3 IT). Despite similar numbers of DEGs after SD, functional enrichment analysis shows a higher level of KEGG, MP and BP specificity for DEGs on L 4/5 IT (Figure 4B), relative to L2/3 IT (Figure 4C). Only in L4/5 IT glutamatergic neurons (ROR-beta positive), SD upregulates certain components of the PI3K, Akt, Ras and MAPK signaling pathways (Figure 4B, Cluster 1 in red, KEGG), such as *Gadd45a*, *Reln*, *p27* and *Sgk1*.

**Figure 4.**
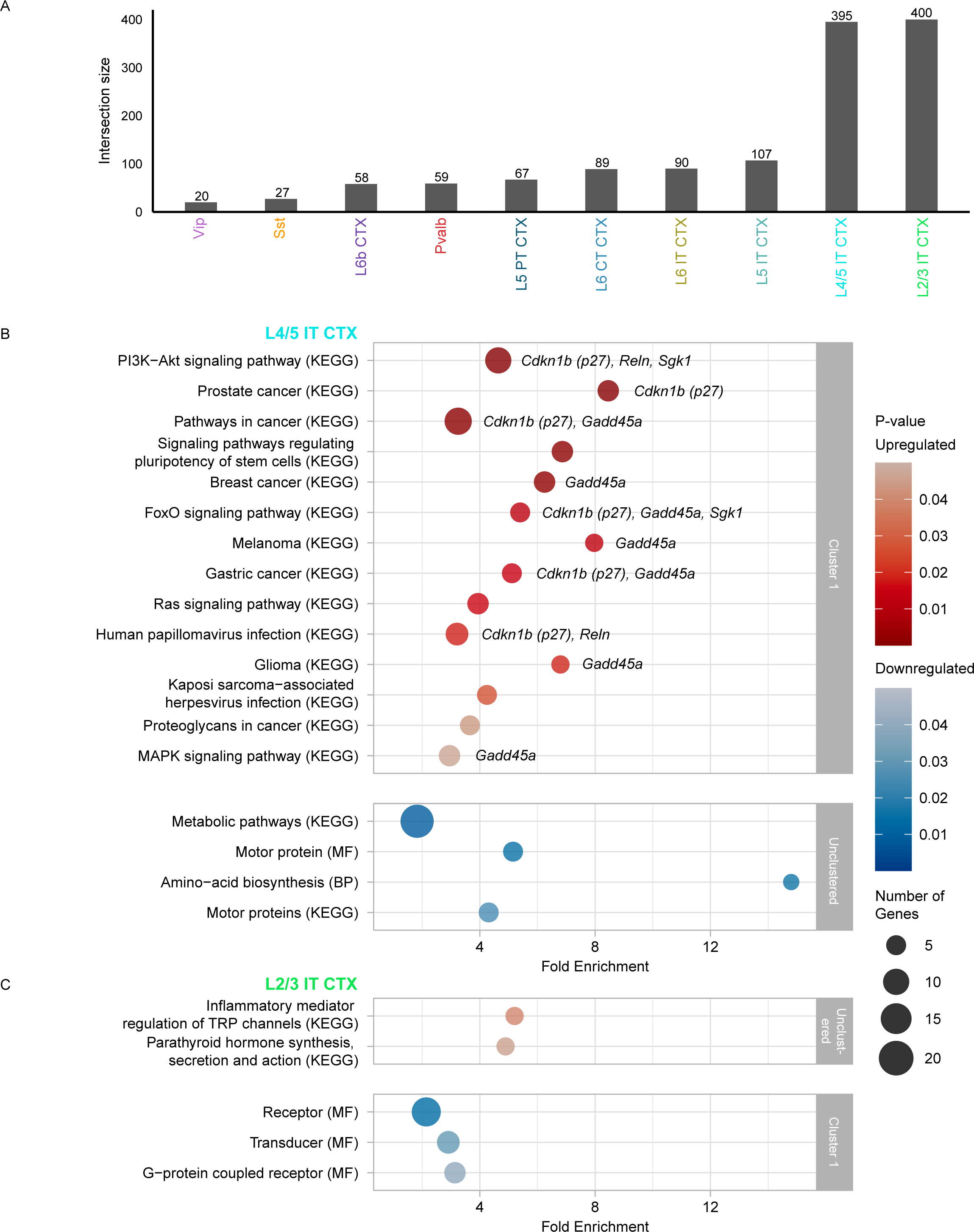
L4/5 IT CTX is preferentially affected by SD relative to other cell-types. **A)** Upset plot showing the unique differentially expressed genes (DEGs) within neuronal cell-types (FDR < 0.05, > 500 nuclei). **B-C)** Bubbles show enriched terms (modified Fisher’s Exact p-value < 0.05) of upregulated (red) and downregulated (blue) DEGs and unique to **B)** L4/5 IT CTX and **C)** L2/3 IT CTX, as compared with the expressed genes within those cell-types. Bubble size corresponds to the number of genes per term (minimum of 3). Gray boxes outline clustered terms (similarity threshold > 0.2, and enrichment score > 1.5). Enrichment scores for each cluster; L4/5 IT CTX Cluster 1 Up (2.17), L2/3 IT CTX Cluster 1 Down (1.59). N=3 per condition. SD, Sleep Deprivation. BP, Uniprot biological process. MF, Uniprot molecular function. KEGG, Kegg pathways.

### Sleep deprivation affects half of the cortical transcriptome and elicits extensive differential isoform usage

Although the effect of SD on brain gene expression is well-documented, it has never been investigated at the isoform level. We chose to perform isoform-level analyses in bulk RNA-seq, in contrast to 10X Chromium snRNA-seq data which has a 3’ bias and limited our ability to recover all transcripts of a gene. We performed differential transcript and gene expression (DTE and DGE) as well as differential transcript usage (DTU) analysis using nonparametric testing of inferential replicate counts after correcting for unwanted variation^17–19^.

Figure 5 shows that after SD we detect 8,505 DEGs (q-value <0.05, Figure 5A, Extended Data Figure 8C, Figure 5 Supplementary Table 1) of 18,334 expressed genes, including several genes previously shown to be affected by SD (highlighted in Figure 5C). These include 83% of positive control genes (558/671) we previously detected across multiple published studies^14^. We then chose the optimal log2 fold-change threshold based on the balance between recovery of positive control genes, while simultaneously not recovering genes less likely to be differentially expressed after SD from public data (Extended Data Figure 9). This resulted in a log2 fold-change >0.2 for downstream analysis. Following DGE analysis, we investigated DTE. We show that 15,525 transcripts (from 10,439 genes) are differentially expressed after SD (q-value <0.05, Figure 5B, Extended Data Figure 8D, Figure 5 Supplementary Table 1), of which 9,709 are downregulated and 5,816 are upregulated after SD. This indicates that transcript level analysis increases our ability to detect differentially expressed genes. Given the discrepancy in downregulated transcripts, we investigated which genes were shared between or unique to DGE and DTE analyses (Figure 5C and 5D): 3,269 upregulated, and 3,836 downregulated genes were shared between analyses, while an additional 3,117 downregulated genes were detected at the transcript level, but not the gene level. This suggests there is more variation happening at the transcript-level, which is obscured when aggregating to the gene-level. To further explore this we focused on eukaryotic initiation factors, which were previously reported to be repressed by SD and are known to mediate the detrimental effects of SD on learning and memory^13,20^. We show that, several eukaryotic initiation factors were significantly differentially expressed at the transcript level that were not detected at the gene level (Extended Data Figure 8E; q-value < 0.05, |log2FC| > 0.20). Our DTE analysis also shows that several genes have both upregulated and downregulated transcripts after SD (e.g. *Bdnf*). To investigate potential opposing effects on transcripts of the same gene we performed differential transcript usage analysis (DTU), to detect which genes changed the proportion of isoforms expressed after SD. We detected 2,314 transcripts (corresponding 1,575 genes, Figure 6 Supplementary Table 1) with significant changes in usage (q-value < 0.05) in response to SD (Figure 6A). These include transcripts that were upregulated at both the gene and transcript level, but SD changes which transcript is primarily transcribed (e.g. *Homer1*, Figure 6B and 6C, Figure 6 Supplementary Table 2), as well as transcripts in which the gene level analysis obscured transcripts being both up and downregulated (e.g. *Bdnf*, Figure 6D and 6E, Figure 6, Supplementary Table 2). For *Bdnf* in particular, *Bdnf I* (201, somatic) increases in proportion, while *Bdnf VI* (205, dendritic) decreases in proportion. These examples suggest that SD may have an effect on splicing and perhaps RNA-binding and transport of somatic versus dendritic isoforms. To further understand which kinds of processes and pathways were affected by isoform switching after SD, we carried out functional enrichment analysis (Figure 7), which revealed 16 enriched pathways/biological processes with 3 clusters of related pathways (Figure 7 Supplementary Table 1). Genes that undergo isoform usage switching in response to SD are related to RNA binding/splicing (e.g. *Rbmx*), chromatin regulation (e.g. *Hdac3*), and kinases (e.g. *Cm1*). Tutorials to perform DGE, DTE and DTU analysis are available through GitHub and at the following website: https://rissolab.github.io/AtlasCortexSD/.

**Figure 5.**
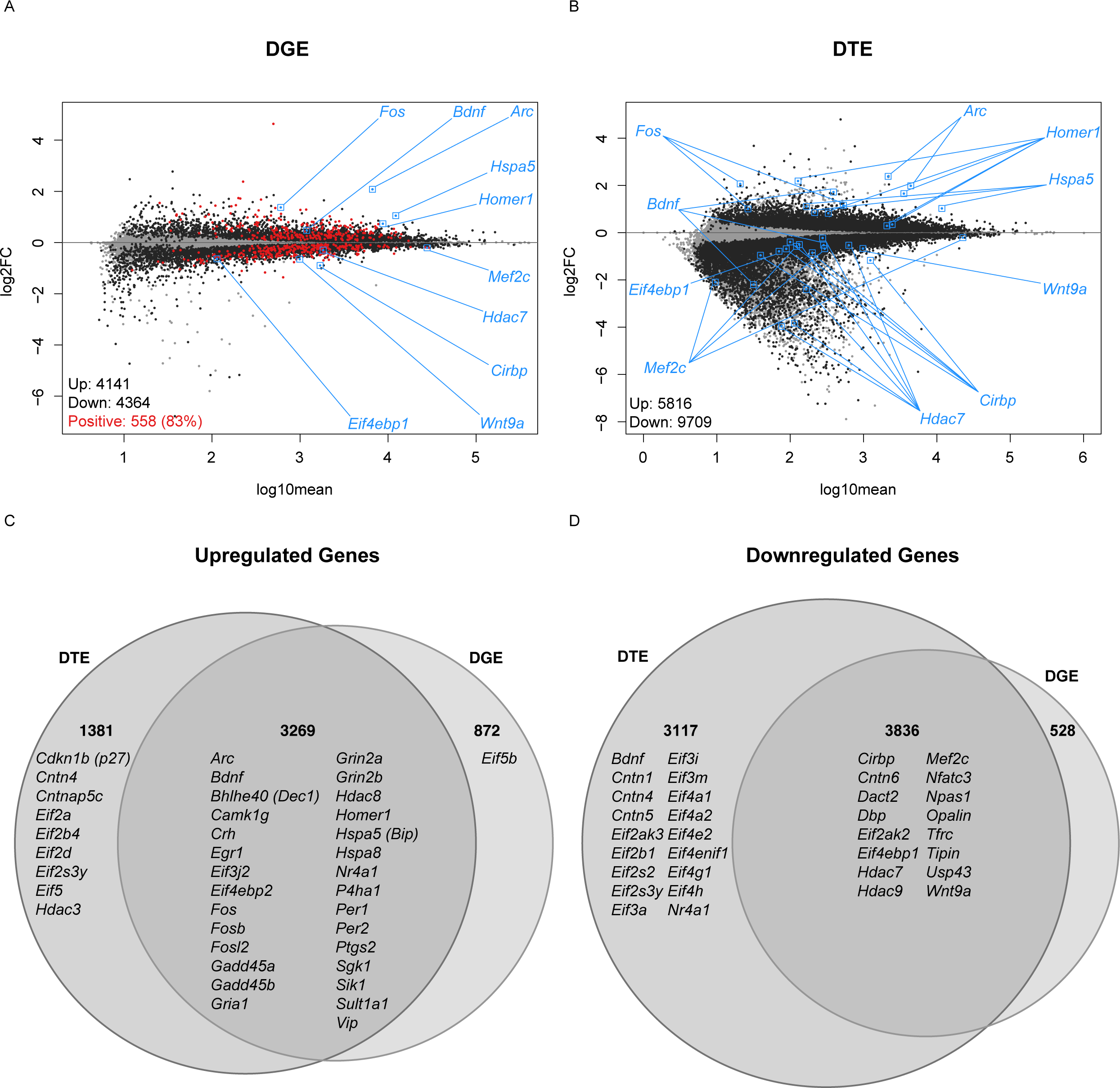
SD predominantly represses transcription at the gene and transcript level. **A)** Differential gene expression (DGE) following SD. On the x-axis, expression is shown as log10mean, and on the y-axis, fold change is shown as log2FC. Expressed genes (18,334) are gray. Significantly differentially expressed genes (8,505), q-value < 0.05, are black. Positive control genes previously shown to respond to sleep deprivation are red^14^. 83.16% positive control genes (558/671) were significantly differentially expressed in response to SD, q-value < 0.05. **B)** Differential transcript expression (DTE) following SD. On the x-axis, expression is shown as log10mean, and on the y-axis, fold change is shown as log2FC. Each gray point shows an expressed transcript (54,030). Significantly differentially expressed transcripts (15,525 from 10,439 genes), q-value < 0.05, are black. A subset of genes from Figure 2 Supplementary Table 2 are shown in **A** and **B**. **C)** Venn diagram shows the intersection of upregulated, significantly differentially expressed genes and upregulated, significantly differentially expressed transcripts. **D)** Venn diagram shows the intersection of downregulated, significantly differentially expressed genes and downregulated, significantly differentially expressed transcripts. Genes from Figure 2 Supplementary Table 2 that have a qvalue < 0.05 and |log2FC| > 0.2 are shown as examples in **C** and **D**. N=5 per condition. SD, Sleep Deprivation.

**Figure 6.**
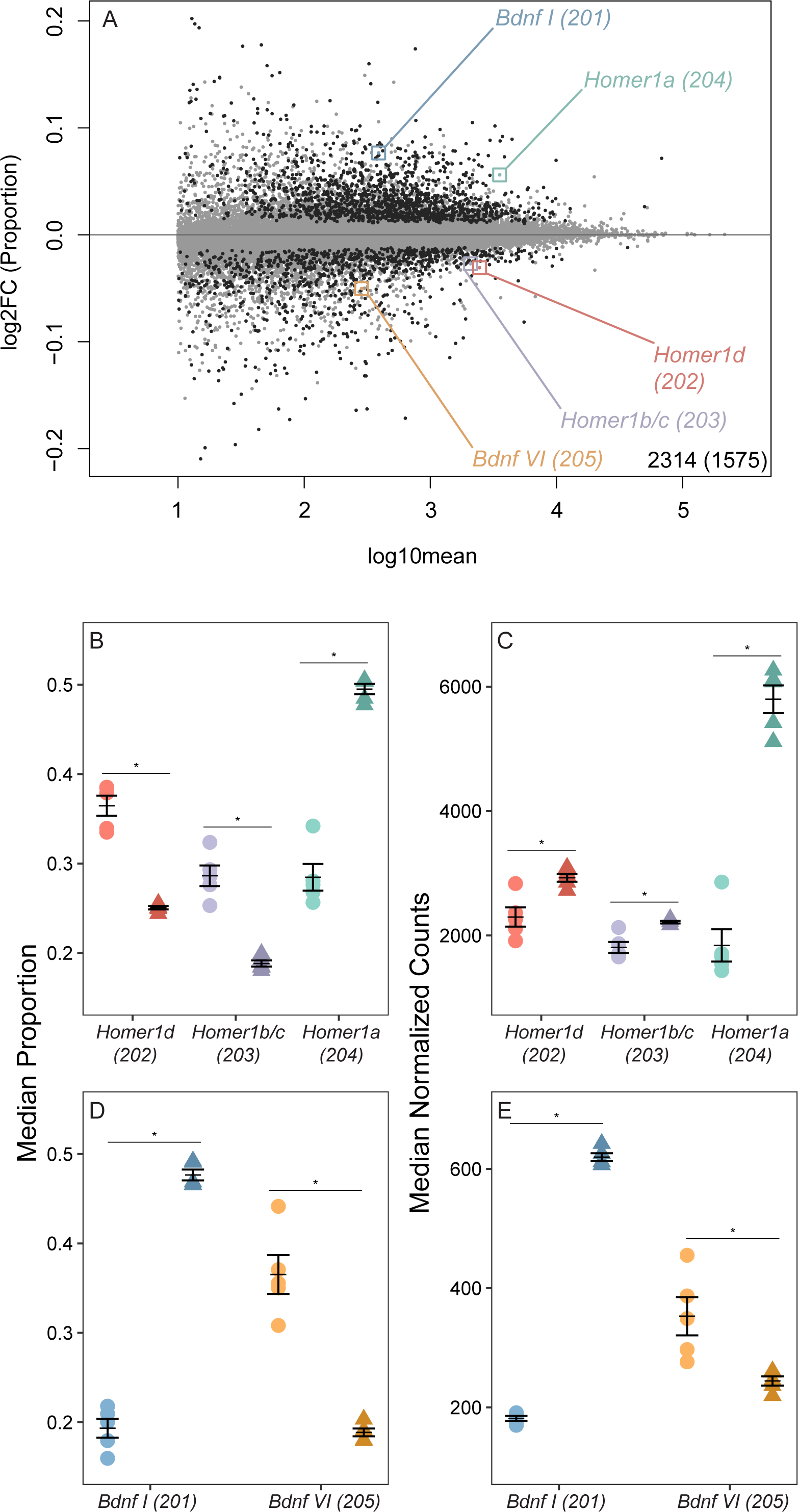
SD promotes differential transcript usage (DTU) for 1,575 genes, including *Homer1* and *Bdnf*. **A)** DTU following SD. On the x-axis, expression is shown as log10mean, and on the y-axis, change in proportion is shown as log2FC. Gray points are expressed transcripts, with a log10mean > 1. In black are transcripts that have significant changes in usage, q-value < 0.05 and log10mean > 1. **B-C)** Dot plot shows the change in **B)** transcript proportion (from 0-1) and **C)** expression levels in normalized transcript counts of “short” (*Homer1a*) and “long” (*Homer1b/c*, *Homer1d*) *Homer1* transcripts^28,29^. **D-E)** Dot plots show the **D)** change in proportion (from 0-1) and **C)** expression levels in normalized transcript counts of “synaptic” (*Bdnf VI*) and “somatic” (*Bdnf I*) *Bdnf* transcripts^30^. For **B-E**, Home cage (HC) animals are circles and SD animals are triangles. Mean ± standard error is shown. N=5 per condition. *Homer1* and *Bdnf* transcripts shown have significant DTE and DTU using Swish, q-value < 0.05^19^. SD, Sleep Deprivation.

**Figure 7.**
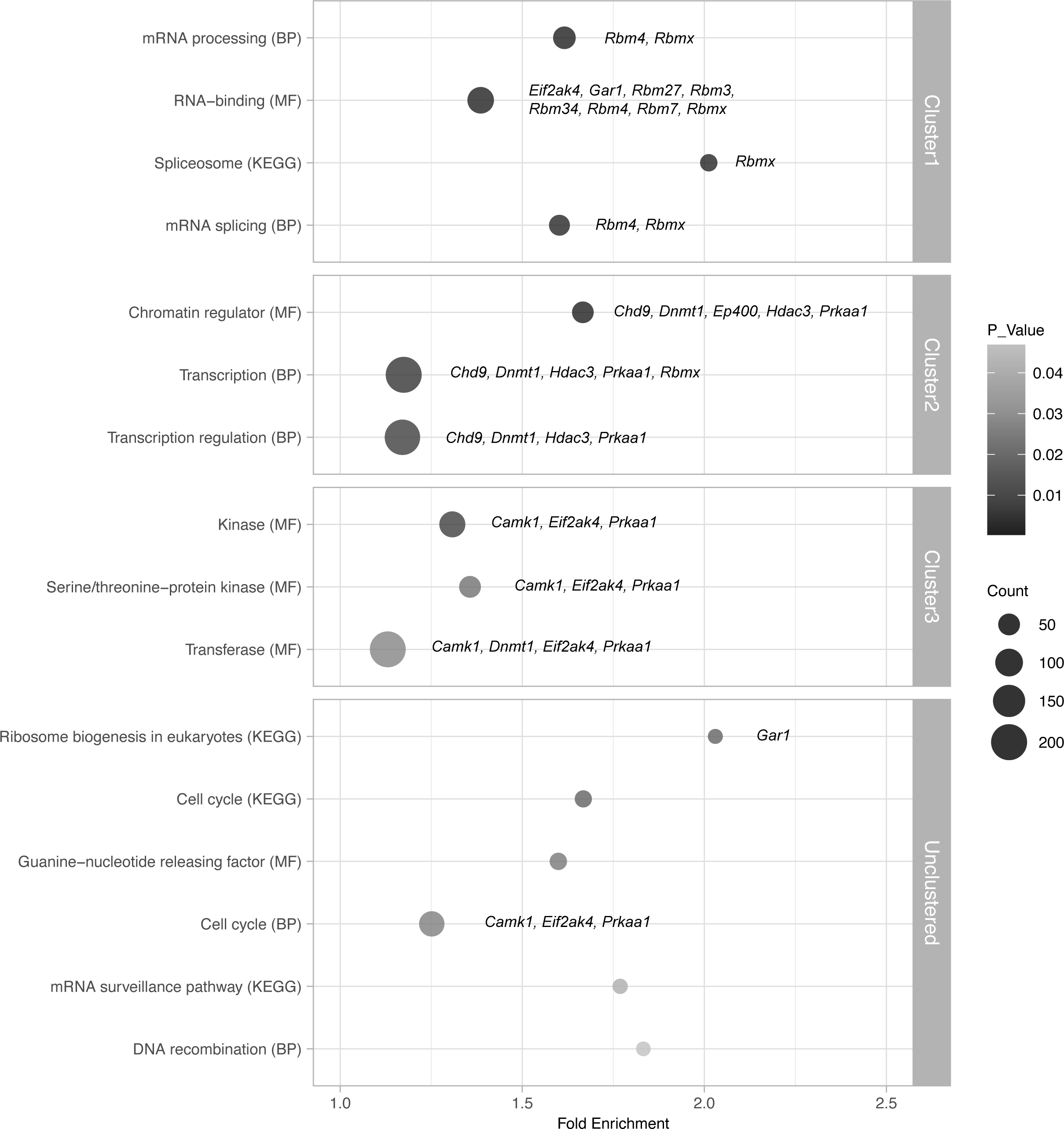
Genes for which SD affects isoform usage are mainly involved in splicing and chromatin regulation. Bubbles show enriched terms (modified Fisher’s Exact p-value < 0.05) following functional enrichment analysis of 1,575 genes that have significant DTU (qvalue < 0.05), as compared with the expressed transcript list. Bubble size corresponds to the number of genes per term (minimum of 3), and color gradient represents p-values. Gray boxes outline clustered terms (similarity threshold > 0.2, and enrichment score > 1.5). Enrichment scores for each cluster; Cluster 1 (3.02), Cluster 2 (2.60), Cluster 3 (1.79). Example genes are shown for each cluster. SD, Sleep Deprivation. DTU, Differential Transcript Usage. BP, Uniprot biological process. MF, Uniprot molecular function. KEGG, KEGG pathways.

## Discussion

In this study we performed for the first time snRNA-seq and bulk RNA-seq in parallel with multiple independent biological replicates in response to sleep deprivation (SD) in the adult male mouse frontal cortex. Prior analyses have focused on bulk gene-level analysis^4–13,21^, or do not include independent biological replicates^22^. Thus to date it was not possible to define what may be occurring at the isoform level or to detect changes specific to particular cell types in response to SD. Because SD has a profound effect on brain function and insufficient sleep is a hallmark of many brain disorders, understanding its molecular impact is not only important to understand the function of sleep but also to understand how behavioral impairments in response to SD arise.

We focused on the frontal cortex because in humans it is the brain area most affected by acute SD^15^. The frontal cortex plays an essential role in higher-order brain processes, including cognition, attention, reward and emotion processing, all of which are affected by lack of sleep. Our snRNA-seq results indicate that SD has a disproportionate effect on neurons (Figure 1). Surprisingly, we do not detect a strong effect of SD on the transcriptome of glia, despite the documented role of glia such as Astrocytes in sleep homeostasis^23^. The low proportion of glia present in our snRNA-seq data (Extended data Figure 1) suggests that to detect the effect of SD in glia it may be necessary to first enrich those populations using glia-specific marker sorting. Even after enrichment, previous transcriptome studies find that only ∼1.4% of the astrocyte transcriptome seems responsive to sleep/wake state^24^. Within neurons, the effect of SD is most prominent in glutamatergic neurons (Figure 2), with over 2,236 genes exclusively regulated in this neuronal type (Figure 3A). Pathway, molecular function and biological process enrichment analysis show that SD disproportionately affects pathways, molecular functions and biological processes involved in neurogenesis, MAPK-PI3K-Akt signaling, circadian rhythms and development in glutamatergic neurons (Figure 3C). Interestingly, down regulation of neurotransmitter receptors by SD is detected in both glutamatergic and GABAergic neurons (KEGG pathway: Neuroactive-ligand receptor interaction, *Vip*, *Sst*, *Gria4*, *Grin2d*). Nonetheless, the disproportionate effect of SD in glutamatergic versus GABAergic neurons may suggest that by altering predominantly the glutamatergic transcriptome SD may alter excitatory/inhibitory balance. Indeed in the visual cortex excitation and inhibition have been shown to be modulated in a sleep-dependent manner in adult mice^25^.

Although is not clear whether or not the rodent frontal cortex possesses a layer 4 as defined in sensory cortices^26^, our data indicates that ROR-beta positive L4/5 IT glutamatergic neurons are the second most abundant type of neuron in the rodent frontal cortex (Figure 2A) and disproportionately responsive to SD at the transcriptome level (Figure 2B). This includes 522 upregulated and 970 downregulated genes after SD (Figure 2A and 2C), 395 of which are unique to these neurons (Figure 4A). ROR-beta expression is more prominent in frontal brain areas in rodents and primates, and drives the development of cortico-thalamic connectivity^27^. In addition to its role as a genetic marker for glutamatergic neurons of layers 4 and 5 in the cortex, ROR-beta is a key transcription factor controlling brain development and differentiation. In ROR-beta positive neurons, SD uniquely upregulates genes (*Reln*, *p27* and *Gadd45a*) and signaling pathways (PI3K, Akt, Ras and MAPK) involved in brain development and neurogenesis and differentiation. The fact that SD specifically affects ROR-beta expressing glutamatergic neurons in the frontal cortex may reflect the importance of sleep in regulating the function of these neurons and thalamocortical circuitry through these pathways.

Using bulk RNA-seq we detect 8,505 differentially expressed genes after SD, while simultaneously recovering 83% of known positive control genes (Figure 5). Although this is not the first study to use bulk RNA-seq to understand the effect of SD on the bulk cortical transcriptome, we detect between 2000-7000 more differentially expressed genes driven by sleep/wake than previously published studies in the cortex and hippocampus^5,6,9,21^. This increase in sensitivity is largely due to differences in methodology on RNA-seq data analysis, such as additional normalization to removal unwanted variance (e.g. RUV-seq) and simultaneous transcript and gene level analysis using nonparametric testing and inferential uncertainty (Fishpond/Swish) ^19^. Incorporating inferential uncertainty alone increases the number of DEGs considerably in our own data compared to our recently published study^9^, while also allowing for differential transcript expression and usage analysis after SD for the first time. We detect 15,525 differentially expressed (Figure 5) and 2,314 differentially used (corresponding 1,575 genes) transcripts (Figure 6A). The latter indicates that SD can induce a large number of isoform switches. Isoform switching is a phenomenon in which the relative contribution of one or many isoforms to the total expression of a particular gene changes significantly between conditions. Two notable examples of the effect of SD on isoform switching are on *Bdnf* and *Homer1* (Figure 6B and C), two genes with known roles in brain function which we show are altered in all neuronal cell-types after SD (Figure 3A). Although the upregulation of *Homer1a* after SD is well-known and commonly referred to as a core molecular correlate of sleep loss^8^, we show that upregulation of *Homer1a* after SD comes at the expense of a lower proportion of long isoforms of *Homer1*, which are more stably bound to the synapse^28,29^. In our analysis, we also find that *Bdnf I* (201, somatic) increases in proportion, while *Bdnf VI* (205, dendritic) decreases in proportion^30^. These examples suggest that SD may have an effect on splicing or RNA-binding. Those processes were not identified as enriched in our snRNA-seq analysis. This may be because only some isoforms of such genes are affected and thus when analyzing differential expression only at the gene level the effect is not detectable or averaged out. Our functional enrichment analysis of genes that are affected by isoform switching after SD using bulk-RNA-seq shows that, indeed, genes involved in splicing and RNA binding are indeed affected by SD at the isoform level (Figure 7). If SD affects splicing and RNA binding, it may perhaps also affect transport of those isoforms to different neuronal compartments (soma vs. synapse). Our results suggest that the role of sleep and SD on isoform expression and transport, and its implication on brain function, needs to be further explored.

Limitations of our study include the fact that we focused on the adult male frontal cortex in our experiments. Future studies aimed at understanding the effect of both sex and developmental age on the transcriptional response to SD with cell type resolution are needed in different brain regions. Recent spatial-transcriptomic studies suggest that different brain regions may have different responses to SD^31^. However, to achieve a full picture of the effect of SD in different brain areas, it will be necessary to combine different technologies (spatial, single cell and bulk RNA sequencing) to balance their strengths and weaknesses. For example, our present study shows that the number of detected genes in snRNA-seq data for most cell-types (Figure 2A) is lower than for bulk RNA-seq. This is because the number of genes detected scales with the number of nuclei sequenced, and even over 15,000 nuclei fall short from detecting the 18,334 expressed genes we obtain using bulk RNA-seq. Thus cell-type resolution may come at a cost of lower sensitivity for differential gene expression. Sensitivity of spatial transcriptomic experiments for differential expression analysis is likely lower than snRNA-seq. Given the inherent differences in the ability to recover DEGs by different technologies, we need to be cautious of the potential for false negatives to alter our interpretation if using only one technology.

Overall, we present the first global transcriptional atlas of the homeostatic response to sleep deprivation in the adult male mouse frontal cortex, combining the advantages of both snRNA-seq and bulk RNA-seq with robust and reproducible data analysis pipelines. We show that SD has a large mostly repressive effect on the cortical transcriptome, that this effect is more prominent in glutamatergic neurons, in particular in L4/5 IT ROR-beta positive neurons. We also show that SD can cause isoform switching of thousands of transcripts. Because sleep and sleep loss are often confounded in rodent studies of brain and behavior, these effects need to be accounted for in *in vivo* transcriptomic studies. As a resource to the neuroscience community, we provide detailed lists of genes, cells and pathways affected by SD as well as tutorials to reproduce our data analysis. Importantly, we made our analyses completely reproducible by sharing all the code used to generate the results of this article and by providing a docker image to run the code with the exact software setup used for this study. In addition, our tutorials can serve as a starting point for the analysis of bulk and snRNA-seq data generated by future studies.

## Methods

### Experimental model and sample details

#### Animals

Wild-type (WT) C57BL6/J mice were housed in standard cages at 24 ± 1°C on a 12:12 h light:dark cycle with food and water *ad libitum*. All experimental procedures were approved by the Institutional Animal Care and Use Committee of Washington State University and conducted in accordance with National Research Council guidelines and regulations for experiments in live animals.

### Single nuclear RNA-seq study after SD

#### Tissue collection

Adult male (8 – 10-week-old) WT C57BL6/J mice were divided into 2 groups (n=3 independent animals per group): sleep deprived (SD5) and home cage controls (HC5). All mice were individually housed. Home cage control mice were left undisturbed and sacrificed 5 h after light onset (ZT5). Mice in the sleep-deprived group were sleep-deprived for 5 h via gentle handling starting at light onset and then sacrificed upon completion of sleep deprivation (ZT5) without allowing for recovery sleep. Mice were sacrificed by live cervical dislocation (alternating between home cage controls and sleep-deprived mice), decapitated, and the frontal cortex was swiftly dissected on a cold block. Tissue was flash frozen in liquid nitrogen and stored at −80°C until processing^9,32^.

#### RNA isolation, library preparation and sequencing

Nuclei were extracted from mouse frontal cortical tissue using the Nuclei PURE Prep kit (NUC201-1KT, Millipore Sigma, Burlington MA, USA) with volumes reduced to a quarter of the recommended amount. Briefly, frontal cortex tissue (∼0.035 g per sample/mouse) were lysed using 2mL glass dounce homogenizers (Kimble-Chase, Vineland, NJ USA) in cold phosphate buffered saline and RNAase inhibitor (03335399001, Roche, Basel, Switzerland). Nuclei was then isolated using a sucrose gradient and ultracentrifugation at 13,000 rcf for 45 mins at 4°C (Sorvall WX-100, F65L-6 x 13.5 rotor, Thermo Fisher, Waltham, MA USA). Isolated nuclei were resuspended in Nuclei PURE Prep kit storage buffer and RNAase inhibitor. Nuclei count and integrity was assessed prior to library preparation.

Single nuclear RNA-seq libraries were generated using the Chromium Single Cell 3’ Solution microfluidics platform (10X Technologies, Pleasonton, CA USA). Single-nuclei libraries were generated from the nuclei suspensions using the 10x Genomics Chromium Controller and Single Cell 3’ Reagent Kits v3 Chemistry following manufacturer’s instructions. Briefly, we targeted capture of 10,000 single nuclei through generation of gel beads in emulsion (GEMs) which allowed partitioning of an individual nuclei along with a bead containing barcoded oligonucleotides. Reverse transcription and barcoding occurred within this emulsion resulting in transcripts from an individual nucleus having a unique molecular identifier (UMI). After barcoding, the emulsion was broken, and the cDNA processed in bulk. The barcoded cDNA was first amplified to generate sufficient mass for library construction and then sample index, P5 and P7 adapters were added for Illumina sequencing. The sizes of 10X cDNA libraries were assessed by Fragment Analyzer with the High Sensitivity NGS Fragment Analysis Kit. The concentrations were measured by StepOnePlus Real-Time PCR System (ThermoFisher Scientific, San Jose, CA) with the KAPA Library Quantification Kit (Kapabiosystems, Wilmington, MA). The libraries were diluted to 2 nM with RSB (10 mM Tris-HCl, pH8.5) and denatured with 0.1 N NaOH. Three pM libraries were loaded onto NextSeq 500 (Illumina, San Diego, CA) for sequencing using the NextSeq 500/550 High Output Kit v2.5 (150 Cycles). The libraries were sequenced from both ends with 28+8+0+91 cycles (read length 100 bp) at an average depth of 40 million paired-end reads per sample.

#### Quantification of raw sequencing reads

We processed the raw sequencing reads in FASTQ files using the Salmon (v. 1.3.0) package using the ‘mapping mode’ that runs in two phases: (i) the indexing step and (ii) the quantification step^33^. To prepare to create the index, we downloaded Gencode (release M25) reference genome (‘GRCm38.primary_assembly.genome.fa.gz’) and reference transcriptome (‘gencode.vM25.transcripts.fa.gz)’, along with GTF coordinates (‘gencode.vM25.annotation.gtf.gz’). To improve the accuracy of quantification estimates from Salmon, we built an index that incorporated a set of genome targets as decoys (https://combine-lab.github.io/alevin-tutorial/2019/selective-alignment). Using the concatenated list of transcriptome targets along with genome targets, we used ‘salmon index’ function to build the index with the flags: ‘--gencodè, ‘--threads 4’, and ‘-k 31’. Next, we used ‘salmon quant’ to perform quantification with the flags ‘--threads 6’ and ‘--numBootstraps 30’. Using this index, we used ‘salmon alevin’ to quantify reads to the gene level with flags ‘--chromiumV3’, ‘--threads 6’, ‘--forceCells 10000’, ‘--dumpFeatures--dumpBfh’, and ‘--numCellBootstraps 30’^34,35^. Next, we created a R/Bioconductor SingleCellExperiment object^36^ with the tximeta (v. 1.15.2) R/Bioconductor package^37^, where we quantified counts for both spliced mRNA and introns using the ‘getFeatureRanges()’ function from the eisaR R/Bioconductor package^38,39^. Also, we used the alevinQC R/Bioconductor package to calculate QC metrics for each sample processed with ‘salmon alevin’.

#### Data setting, quality control, normalization and doublets removal

We summed the UMI counts of spliced mRNA and introns sharing the same Ensembl Gene ID. To identify mitochondrial genes, we retrieved the chromosome location of each Ensembl Gene ID with the EnsDb.Mmusculus.v79 (v. 2.99.0) R/Bioconductor package^40^. We then split the data into six SingleCellExperiment objects, one for each mouse.

For each sample, we used the scuttle (v. 1.8.4) R/Bioconductor package to detect low-quality and damaged droplets^41^. Particularly, we computed per-cell quality-control metrics with the *perCellQCMetrics* function; these metrics include the sum of UMI counts, the number of detected genes, and the percentage of mitochondrial counts. Lastly, for each sample, we removed potential doublets with the scDblFinder (v. 1.12.0) R/Bioconductor package^42^, using the *computeDoubletDensity* function to calculate the scores and the *doubletThresholding* function to set the doublet scores threshold with the *griffiths* method.

Overall, our quality control procedure retained 52,651 high-quality nuclei, with an average of 8,775 nuclei per mouse.

### Cell-type annotation and validation of cell-type labels

To identify cell types, we used the Allen Whole Cortex & Hippocampus - 10x genomics data http://portal.brain-map.org/atlases-and-data/rnaseq/mouse-whole-cortex-and-hippocampus-10x) as a reference dataset^16^. This dataset was imported with the *AllenInstituteBrainData* function of the AllenInstituteBrainData (v. 0.99.1) package (available at https://github.com/drighelli/AllenInstituteBrainData). We then selected the “Non-Neuronal”, “Glutamatergic’’ and “GABAergic” clusters coming from the Visual Cortex (VIS, VISl, VISm, VISp) to annotate our dataset. For computational issues, we selected a random subset of 100,000 cortical cells.

To identify the best cell annotation method, we used two datasets of primary motor cortex tissue. The first dataset, “10x Nuclei v3 Broad,” (http://data.nemoarchive.org/biccn/lab/zeng/transcriptome/sncell/10x_v3/mouse/processed/anal <u>ysis/10X_nuclei_v3_Broad/</u>) was from the Broad (Macosko Lab), while the second dataset, “10X Nuclei v2 AIBS,” (http://data.nemoarchive.org/biccn/lab/zeng/transcriptome/sncell/10x_v2/mouse/processed/anal <u>ysis/10X_nuclei_v2_AIBS/</u>) was from the Allen Institute for Brain Science ^16^.

We then annotated these datasets using two methods: Azimuth and SingleR. For Azimuth, the reference data was converted into a Seurat object and into an Azimuth compatible object, using the *AzimuthReference* function of the Azimuth (v. 0.4.6) package https://satijalab.github.io/azimuth/articles/run_azimuth_tutorial.html^43^. Then query samples were merged and converted into a Seurat object. Cell annotation was computed using the *RunAzimuth* function of the Azimuth package. The t-SNE and the UMAP embeddings were computed using the *RunTSNE* and *RunUMAP* functions of the Seurat (v. 4.3.0) package ^44^, https://cran.r-project.org/web/packages/Seurat/index.html) with *seed.use = 1*. For SingleR, the reference dataset was aggregated across groups of cell types and was normalized, using the *aggregateAcrossCells* and the *logNormCounts* functions of the scuttle (v. 1.8.4) package. Then, cell annotation was computed using the *SingleR* function of the SingleR (v. 2.0.0) R/Bioconductor package ^45^, https://bioconductor.org/packages/release/bioc/html/SingleR.html). We found that Azimuth was the best-performing method on these already annotated datasets and hence we chose this annotation method for the annotation of our rodent PFC snRNA-seq dataset. In addition, to evaluate the cell-type assignments in our dataset, we visualized cell-type specific markers based on references ^46–50^, with a heatmap of the log-normalized count average in each group. We used the *pheatmap* function of the pheatmap (v. 1.0.12) package (https://cran.r-project.org/web/packages/pheatmap/index.html). As an additional quality control, we checked if there were cell types with a low proportion of intronic reads, as this could be a sign of cytoplasmic RNA (likely from cell debris) and assigned incorrectly to nuclei. All cell types had a high proportion of intronic reads, as expected in single-nuclear RNA-seq^51^.

To visualize the assigned cell-type labels in two dimensions, the UMAP embeddings were computed using the *DimPlot* function of the Seurat package, with option *reduction = “integrated_dr”*, where “integrated_dr” is the supervised principal component analysis obtained by the Azimuth method. Finally, a pseudo-bulk level Multidimensional Scaling (MDS) plot was created with the *pbMDS* function of the muscat (v. 1.12.1) R/Bioconductor package^52^. Each point represents one subpopulation-sample instance; points are colored by subpopulation and shaped by treatment.

### Pseudo-bulk differential expression analysis for snRNA-seq data

For each neuronal cell type with more than 500 nuclei, differential gene expression analysis was carried out with a negative binomial generalized linear model (GLM) on pseudo-bulk samples. Specifically, we created the pseudo-bulk samples with the function *aggregateAcrossCells* of the scuttle package by summing the counts of each gene for each cell type and mouse combination.

Then, we normalized the raw counts for each cell type with the upper-quartile method, using the *betweenLaneNormalization* function of the EDASeq (v. 2.32.0) R/Bioconductor package with option *which=“upper”*^53^. To account for latent confounders, we computed the factors of unwanted variation on the normalized data, using the *RUVs* function of the RUVSeq R/Bioconductor (v. 1.32.0) package with *k=2*^17,18^, and using a list of genes previously characterized as non-differential in sleep deprivation in a large microarray meta-analysis^14^, herein referred to as “negative control genes.” Specifically, 10% of negative control genes were randomly selected to be used for evaluation and the remaining control genes were used to fit RUV normalization.

We then used the edgeR R/Bioconductor (v. 3.40.2) package to perform differential expression after filtering the lowly expressed genes with the *filterByExpr* function (with parameters: *min.count = 10, min.total.count = 15, large.n = 10, min.prop = 0.7)*^54^. The raw counts were normalized with the upper-quartile method, using the function *calcNormFactors*^55^. The factors of unwanted variation were added to the design matrix. The differential gene expression analysis was performed with the function *glmLRT* by specifying “SD-HC” (Sleep Deprived vs Home Cage Control) as contrast. We used the Benjamini-Hochberg procedure to control for the false discovery rate (FDR), i.e., we considered as differentially expressed those genes that had an adjusted p-value less than 5%^56^.

For each cell type, we visualized differentially expressed genes (DEGs) with volcano plots and assessed the model’s goodness-of-fit by visualizing the p-value histograms. We incorporated cross-study, cross-brain tissue positive controls (Additional File 2 from Gerstner et al., 2016 to evaluate the performance of our differential gene expression pipeline.

For glutamatergic and GABAergic neurons, we used the *upset* function of the UpSetR (v. 1.4.0) package to compare the lists of differentially expressed genes within each cell type with more than 500 nuclei^57^.

To better understand which genes were shared between glutamatergic and GABAergic cell types, the union of all glutamatergic DEGs and GABAergic DEGs was determined. To visualize shared and unique genes, a Venn Diagram was generated using the *venn.diagram* function of the VennDiagram package (v. 1.7.3, https://cran.r-project.org/web/packages/VennDiagram/index.html). Select genes were highlighted.

### Functional enrichment analysis of snRNA-seq data

Additionally, functional enrichment analysis of genes that were shared between glutamatergic and GABAergic cell types, or unique to a given category (glutamatergic only or GABAergic only) were subjected to functional annotation using the Database for Annotation, Visualization and Integrated Discovery v2021 (DAVID)^58,59^. Prior to the analysis, genes were separated by fold change to obtain one list of upregulated and one list of downregulated genes per category.

Genes that were upregulated in one cell type, but downregulated in another were excluded from analysis. The following categories were used: Uniprot Biological Process, Uniprot Molecular Function (https://www.uniprot.org) and KEGG Pathways (https://www.genome.jp/kegg/pathway.html). Enrichment was relative to the union of all expressed genes within a category: unique to glutamatergic cell types or unique to GABAergic cell types. An EASE Score < 0.05 and similarity threshold > 0.20 were used to allow for inclusive clustering. Clustered and unclustered terms were visualized with a bubble plot using the *ggplot* function from the ggplot2 (v. 3.4.2, https://cran.r-project.org/web/packages/ggplot2/index.html) package, with the size of the bubbles corresponding to the number of genes within a term. For glutamatergic and GABAergic bubble plots, clustered terms were reduced to one bubble, with the size of the bubble corresponding to the union of genes within all terms in that category. Duplicated genes were removed. Fold enrichment is visualized along the x-axis. For clustered terms, the geometric mean of the fold enrichments was determined and plotted along the x-axis. P-values are shown as a color gradient, red for upregulated and blue for downregulated. For clustered terms, the geometric mean of the p-values was plotted.

Functional enrichment of genes that were differentially expressed, and unique to a cell type, was performed using DAVID. DGE lists were separated by fold change to obtain one list of upregulated, and one list of downregulated genes per cell type. The same categories were used as detailed above: Uniprot Biological Process, Uniprot Molecular Function and KEGG Pathways. Again, an EASE Score < 0.05 and similarity threshold > 0.20 were used to allow for inclusive clustering. For L2/3 IT CTX and L4/5 IT CTX, clustered and unclustered terms were visualized with a bubble plot using the *ggplot* function from the ggplot2 (v. 3.4.2, https://cran.r-project.org/web/packages/ggplot2/index.html) package, with the size of the bubbles corresponding to the number of genes within a term. The fold enrichment is visualized along the x-axis. P-values are shown as a color gradient, red for upregulated and blue for downregulated.

### Bulk RNA-seq gene expression study after SD

#### Tissue collection

Adult male (8 – 10-week-old) WT C57BL6/J mice were divided into 2 groups (n=5 independent animals per group): sleep deprived (SD5) and home cage controls (HC5). All mice were individually housed. Home cage control mice were left undisturbed and sacrificed 5 h after light onset (ZT5). Mice in the sleep deprived group were sleep deprived for 5 h via gentle handling starting at light onset and then sacrificed upon completion of sleep deprivation (ZT5) without allowing for recovery sleep. Mice were sacrificed by live cervical dislocation (alternating between home cage controls and sleep deprived mice), decapitated, and the frontal cortex was swiftly dissected on a cold block. Tissue was flash frozen in liquid nitrogen and stored at −80°C until processing ^9,32^. This protocol was repeated over a 5 day period, and all tissue was collected within the first 15 min of the hour.

#### RNA isolation, library preparation and sequencing

Frontal cortex tissue was homogenized in Qiazol buffer (Qiagen, Hilden, Germany) using a TissueLyser (Qiagen) and all RNA was extracted using the Qiagen RNAeasy kit (Qiagen) on the same day. The integrity of total RNA was assessed using Fragment Analyzer (Advanced Analytical Technologies, Inc., Ankeny, IA) with the High Sensitivity RNA Analysis Kit (Advanced Analytical Technologies, Inc.). RNA Quality Numbers (RQNs) from 1 to 10 were assigned to each sample to indicate its integrity or quality. “10” stands for a perfect RNA sample without any degradation, whereas “1” marks a completely degraded sample. RNA samples with RQNs ranging from 8 to 10 were used for RNA library preparation with the TruSeq Stranded mRNA Library Prep Kit (Illumina, San Diego, CA). Briefly, mRNA was isolated from 2.5 µg of total RNA using poly-T oligo attached to magnetic beads and then subjected to fragmentation, followed by cDNA synthesis, dA-tailing, adaptor ligation and PCR enrichment. The sizes of RNA libraries were assessed by Fragment Analyzer with the High Sensitivity NGS Fragment Analysis Kit (Advanced Analytical Technologies, Inc.). The concentrations of RNA libraries were measured by StepOnePlus Real-Time PCR System (ThermoFisher Scientific, San Jose, CA) with the KAPA Library Quantification Kit (Kapabiosystems, Wilmington, MA). The libraries were diluted to 2 nM with Tris buffer (10 mM Tris-HCl, pH8.5) and denatured with 0.1 N NaOH. Eighteen pM libraries were clustered in a high-output flow cell using HiSeq Cluster Kit v4 on a cBot (Illumina). After cluster generation, the flow cell was loaded onto HiSeq 2500 for sequencing using HiSeq SBS kit v4 (Illumina). DNA was sequenced from both ends (paired-end) with a read length of 100 bp. The average depth for all samples was 52 million read pairs.

#### Quantification of raw sequencing reads

To process the raw sequencing reads from the snRNA-seq experiments, we again used Salmon along with the same index built with decoys, which is detailed in the ‘Quantification of raw sequencing reads’ in the ‘Single nuclear RNA-seq study after SD’ section. Again, we used the tximeta to create a SummarizedExperiment object at the transcript level and a second object summarized to the gene level using the ‘summarizeToGene()’ from tximeta.

#### Differential Gene and Transcript Expression

Following tximeta, inferential replicates were scaled and filtered using the default parameters in the fishpond R/Bioconductor package (v. 2.4.0) so that only features with a minimum of 3 samples with a minimum count of 10 reads remained, herein referred to as “expressed” ^19^. To correct for unwanted variation, raw counts were first normalized with the upper-quartile method, using the *betweenLaneNormalization* function of the EDASeq (v. 2.32.0) R/Bioconductor package with option *which=“upper”* ^53^. Next, *RUVs* (k=4) from the RUVseq (v. 1.32.0) R/Bioconductor package was used to generate ‘W,’ the factors of unwanted variation^17,18^. For RUVs at the gene level, we implemented the same list of genes less likely to be affected by sleep deprivation according to microarrays detailed previously, herein referred to as “negative control genes”^14^. For RUVs at the transcript level, we used all expressed transcripts as controls. We then used ‘removeBatchEffect’ from the limma R/Bioconductor package (v. 3.54.0) to remove the variation from the inferential replicates^60^. After correcting for unwanted variation, differential expression was performed using *Swish* from the fishpond package. Genes and transcripts with a q-value < 0.05 (multiple test corrected p-value) were deemed to be significantly differentially expressed. We incorporated the same cross-study, cross-brain tissue positive controls detailed previously (Additional File 2 from ^14^ to evaluate the performance of our differential gene expression pipeline. We recovered 83.2% (558/671) of the positive control genes detected in the matrix at the gene level. To determine which genes were only detected with expression analysis at the gene or transcript level, the *venn.diagram* function of the VennDiagram package (v. 1.7.3, https://cran.r-project.org/web/packages/VennDiagram/index.html) was implemented.

#### Differential Transcript Usage

During our differential expression analysis, we discovered genes that had both upregulated and downregulated transcripts. Therefore, we decided to perform differential transcript usage (DTU) analysis, to detect which genes had transcripts with differential proportion in response to sleep deprivation. To do so, we incorporated ‘isoformProportions’ from the fishpond package, to convert the counts of inferential replicates to proportions before proceeding with *Swish*. To increase the reproducibility of the results presented, a secondary filter was immediately applied following the initial filtering which kept transcripts with a minimum of 10 reads across 3 samples. Only transcripts that had a log10mean > 1 were kept, removing transcripts with low counts that passed the initial filtering.

To better visualize the change in the proportion of transcripts within genes of particular interest (*Homer1* and *Bdnf*), dot plots were generated using the *ggplot* function from the ggplot2 (v. 3.4.2, https://cran.r-project.org/web/packages/ggplot2/index.html) package. Briefly, for each biological replicate, the median of the inferential replicates was determined to obtain one value per transcript per animal. Transcripts were only included if they had significant changes in both proportion and expression, q-value < 0.05. Additionally, the mean and standard error within each condition was determined for each transcript and are shown.

In addition to plotting the proportions of transcripts that had significant changes in proportion, dot plots were also generated to show the normalized counts of the same transcripts following DTE analysis. To generate these plots, the median of the inferential replicates was determined following swish to obtain one value per transcript per animal. The normalized counts were only plotted for transcripts that had significant changes in proportion and expression, q-value < 0.05. Additionally, the mean and standard error within each condition was determined for each transcript and are shown.

#### Functional Enrichment Analysis of bulkRNA-seq

Functional enrichment analysis of the 1,575 genes with significant differential transcript usage (q-value < 0.05) was performed using DAVID as detailed in the ‘Functional enrichment analysis of snRNA-seq data’ section within the ‘Single nuclear RNA-seq study after SD’. The same categories were used as detailed previously: Uniprot Biological Process, Uniprot Molecular Function and KEGG Pathways. Enrichment was relative to the expressed genes after the initial filter preserving transcripts with a minimum of 10 reads across 3 samples, and before the additional log10mean filter. A p-value threshold for gene enrichment analysis (EASE Score) < 0.05 was used. A similarity threshold > 0.20 was used to allow for inclusive clustering. Both clustered and unclustered terms were visualized with a bubble plot using the *ggplot* function from the ggplot2 (v. 3.4.2, https://cran.r-project.org/web/packages/ggplot2/index.html) package. For functional annotation of genes with significant changes in expression in response to sleep deprivation, please see Muheim et al., 2023.

## Data Availability

Sequencing data have been deposited in NCBI’s Gene Expression Omnibus (GEO) under the accession number <u>GSE211088</u>. The bulk RNA-seq replicates (5 SD, 5 HC) were previously deposited in GEO under accession number <u>GSE113754</u>, and downloaded from GEO for this analysis. Supplementary files can be accessed at Zenodo^61^.

## Code Availability

The code used in this article can be accessed via Github through the following link: https://github.com/PeixotoLab/RNAseq_sleep. The Allen Whole Cortex & Hippocampus - 10x genomics (v2021) reference dataset used for single-nuclear analysis, in a SingleCellExperiment object, has been made available at https://github.com/drighelli/AllenInstituteBrainData. RMarkdown tutorials for reference-based cell-type annotation and differential expression and usage analyses can be found in Supplementary Software and at the following website: https://rissolab.github.io/AtlasCortexSD/index.html.

## Supporting information

Inventory of Supplementary information

## Acknowledgements

This work was supported by the National Institute of General Medical Sciences (NIGMS) under project number R35GM147020 and by the National Institute of Neurological Disorders and Stroke (NINDS) under project number R56NS124805 to L.P. S.C.H. and D. Ris. are also supported by CZF2019-002443 from the Chan Zuckerberg Initiative DAF, an advised fund of Silicon Valley Community Foundation.

## Author Contributions

K.F., E.M., H.S., K.S. and L.P. developed methodology and conducted the investigation. K.F., E.Z., C.M., D.Rig. and S.C.H. adapted and wrote the bioinformatic code, and carried out the statistical analysis. K.F., E.Z., L.P., S.C.H. and D.Ris. drafted the paper. K.F., C.M., M.G.F, S.C.H., D.Ris. and L.P. edited and reviewed the paper. L.P., S.C.H. and D.Ris. supervised and managed the study. L.P. conceptualized the study, provided resources and acquired funding.

## Competing Interests

The authors declare no competing interests.

## Notes

https://rissolab.github.io/AtlasCortexSD

https://doi.org/10.5281/zenodo.10041833

https://www.ncbi.nlm.nih.gov/geo/query/acc.cgi?acc=GSE211088

### Competing Interest Statement

The authors have declared no competing interest.

## References

1. Lyons LC, Vanrobaeys Y, Abel T. Sleep and memory: The impact of sleep deprivation on transcription, translational control, and protein synthesis in the brain. J Neurochem. 2023;166(1):24–46. doi:10.1111/jnc.15787

2. Shen Y, Lv Q kun, Xie W ye, et al. Circadian disruption and sleep disorders in neurodegeneration. Transl Neurodegener. 2023;12(1):8. doi:10.1186/s40035-023-00340-6

3. Veatch OJ, Maxwell-Horn AC, Malow BA. Sleep in Autism Spectrum Disorders. Curr Sleep Med Rep. 2015;1(2):131–140. doi:10.1007/s40675-015-0012-1

4. Cirelli C, Gutierrez CM, Tononi G. Extensive and divergent effects of sleep and wakefulness on brain gene expression. Neuron. 2004;41(1):35–43. doi:10.1016/s0896-6273(03)00814-6

5. Gaine ME, Bahl E, Chatterjee S, Michaelson JJ, Abel T, Lyons LC. Altered hippocampal transcriptome dynamics following sleep deprivation. Mol Brain. 2021;14(1):125. doi:10.1186/s13041-021-00835-1

6. Hor CN, Yeung J, Jan M, et al. Sleep–wake-driven and circadian contributions to daily rhythms in gene expression and chromatin accessibility in the murine cortex. Proc Natl Acad Sci. 2019;116(51):25773–25783. doi:10.1073/pnas.1910590116

7. Mackiewicz M, Shockley KR, Romer MA, et al. Macromolecule biosynthesis: a key function of sleep. Physiol Genomics. 2007;31(3):441–457. doi:10.1152/physiolgenomics.00275.2006

8. Maret S, Dorsaz S, Gurcel L, et al. Homer1a is a core brain molecular correlate of sleep loss. Proc Natl Acad Sci U S A. 2007;104(50):20090–20095. doi:10.1073/pnas.0710131104

9. Muheim CM, Ford K, Medina E, Singletary K, Peixoto L, Frank MG. Ontogenesis of the molecular response to sleep loss. Neurobiol Sleep Circadian Rhythms. 2023;14:100092. doi:10.1016/j.nbscr.2023.100092

10. Naidoo N, Giang W, Galante RJ, Pack AI. Sleep deprivation induces the unfolded protein response in mouse cerebral cortex. J Neurochem. 2005;92(5):1150–1157. doi:10.1111/j.1471-4159.2004.02952.x

11. Noya SB, Colameo D, Brüning F, et al. The forebrain synaptic transcriptome is organized by clocks but its proteome is driven by sleep. Science. 2019;366(6462):eaav2642. doi:10.1126/science.aav2642

12. Terao A, Wisor JP, Peyron C, et al. Gene Expression in the Rat Brain during Sleep Deprivation and Recovery Sleep: An Affymetrix GeneChip® Study. Neuroscience. 2006;137(2):593–605. doi:10.1016/j.neuroscience.2005.08.059

13. Vecsey CG, Peixoto L, Choi JHK, et al. Genomic analysis of sleep deprivation reveals translational regulation in the hippocampus. Physiol Genomics. 2012;44(20):981–991. doi:10.1152/physiolgenomics.00084.2012

14. Gerstner JR, Koberstein JN, Watson AJ, et al. Removal of unwanted variation reveals novel patterns of gene expression linked to sleep homeostasis in murine cortex. BMC Genomics. 2016;17(Suppl 8):727. doi:10.1186/s12864-016-3065-8

15. Verweij IM, Romeijn N, Smit DJ, Piantoni G, Van Someren EJ, van der Werf YD. Sleep deprivation leads to a loss of functional connectivity in frontal brain regions. BMC Neurosci. 2014;15(1):88. doi:10.1186/1471-2202-15-88

16. Yao Z, van Velthoven CTJ, Nguyen TN, et al. A taxonomy of transcriptomic cell types across the isocortex and hippocampal formation. Cell. 2021;184(12):3222–3241.e26. doi:10.1016/j.cell.2021.04.021

17. Peixoto L, Risso D, Poplawski SG, et al. How data analysis affects power, reproducibility and biological insight of RNA-seq studies in complex datasets. Nucleic Acids Res. 2015;43(16):7664-7674. doi:10.1093/nar/gkv736

18. Risso D, Ngai J, Speed TP, Dudoit S. Normalization of RNA-seq data using factor analysis of control genes or samples. Nat Biotechnol. 2014;32(9):896–902. doi:10.1038/nbt.2931

19. Zhu A, Srivastava A, Ibrahim JG, Patro R, Love MI. Nonparametric expression analysis using inferential replicate counts. Nucleic Acids Res. 2019;47(18):e105. doi:10.1093/nar/gkz622

20. Tudor JC, Davis EJ, Peixoto L, et al. Sleep deprivation impairs memory by attenuating mTORC1-dependent protein synthesis. Sci Signal. 2016;9(425):ra41. doi:10.1126/scisignal.aad4949

21. Bjorness TE, Kulkarni A, Rybalchenko V, et al. An essential role for MEF2C in the cortical response to loss of sleep in mice. eLife. 2020;9:e58331. doi:10.7554/eLife.58331

22. Jha PK, Valekunja UK, Ray S, Nollet M, Reddy AB. Single-cell transcriptomics and cell-specific proteomics reveals molecular signatures of sleep. Commun Biol. 2022;5(1):1–16. doi:10.1038/s42003-022-03800-3

23. Ingiosi AM, Frank MG. Goodnight, astrocyte: waking up to astroglial mechanisms in sleep. FEBS J. 2023;290(10):2553–2564. doi:10.1111/febs.16424

24. Bellesi M, de Vivo L, Tononi G, Cirelli C. Effects of sleep and wake on astrocytes: clues from molecular and ultrastructural studies. BMC Biol. 2015;13:66. doi:10.1186/s12915-015-0176-7

25. Bridi MCD, Zong FJ, Min X, et al. Daily Oscillation of the Excitation-Inhibition Balance in Visual Cortical Circuits. Neuron. 2020;105(4):621–629.e4. doi:10.1016/j.neuron.2019.11.011

26. Anastasiades PG, Carter AG. Circuit organization of the rodent medial prefrontal cortex. Trends Neurosci. 2021;44(7):550–563. doi:10.1016/j.tins.2021.03.006

27. Shibata M, Pattabiraman K, Lorente-Galdos B, et al. Regulation of prefrontal patterning and connectivity by retinoic acid. Nature. 2021;598(7881):483-488. doi:10.1038/s41586-021-03953-x

28. Clifton NE, Trent S, Thomas KL, Hall J. Regulation and Function of Activity-Dependent Homer in Synaptic Plasticity. Mol Neuropsychiatry. 2019;5(3):147–161. doi:10.1159/000500267

29. Shiraishi-Yamaguchi Y, Furuichi T. The Homer family proteins. Genome Biol. 2007;8(2):206. doi:10.1186/gb-2007-8-2-206

30. Chiaruttini C, Sonego M, Baj G, Simonato M, Tongiorgi E. BDNF mRNA splice variants display activity-dependent targeting to distinct hippocampal laminae. Mol Cell Neurosci. 2008;37(1):11–19. doi:10.1016/j.mcn.2007.08.011

31. Vanrobaeys Y, Mukherjee U, Langmack L, et al. Mapping the spatial transcriptomic signature of the hippocampus during memory consolidation. Nat Commun. 2023;14(1):6100. doi:10.1038/s41467-023-41715-7

32. Ingiosi A, Schoch H, Wintler TP, et al. Shank3 Modulates Sleep and Expression of Circadian Transcription Factors. Kim E, ed. eLife. 2019;8:e42819. doi:10.7554/eLife.42819

33. Patro R, Duggal G, Love MI, Irizarry RA, Kingsford C. Salmon provides fast and bias-aware quantification of transcript expression. Nat Methods. 2017;14(4):417–419. doi:10.1038/nmeth.4197

34. Srivastava A, Malik L, Smith T, Sudbery I, Patro R. Alevin efficiently estimates accurate gene abundances from dscRNA-seq data. Genome Biol. 2019;20(1):65. doi:10.1186/s13059-019-1670-y

35. Srivastava A, Malik L, Sarkar H, Patro R. A Bayesian framework for inter-cellular information sharing improves dscRNA-seq quantification. Bioinformatics. 2020;36(Suppl 1):i292–i299. doi:10.1093/bioinformatics/btaa450

36. Amezquita RA, Lun ATL, Becht E, et al. Orchestrating Single-Cell Analysis with Bioconductor. Nat Methods. 2020;17(2):137–145. doi:10.1038/s41592-019-0654-x

37. Love MI, Soneson C, Hickey PF, et al. Tximeta: Reference sequence checksums for provenance identification in RNA-seq. PLoS Comput Biol. 2020;16(2):e1007664. doi:10.1371/journal.pcbi.1007664

38. Gaidatzis D, Burger L, Florescu M, Stadler MB. Erratum: Analysis of intronic and exonic reads in RNA-seq data characterizes transcriptional and post-transcriptional regulation. Nat Biotechnol. 2016;34(2):210–210. doi:10.1038/nbt0216-210a

39. Soneson C, Srivastava A, Patro R, Stadler MB. Preprocessing choices affect RNA velocity results for droplet scRNA-seq data. PLoS Comput Biol. 2021;17(1):e1008585. doi:10.1371/journal.pcbi.1008585

40. Rainer, Johannes. EnsDb.Mmusculus.v79. Bioconductor. Published 2017. Accessed November 3, 2023. http://bioconductor.org/packages/EnsDb.Mmusculus.v79/

41. McCarthy DJ, Campbell KR, Lun ATL, Wills QF. Scater: pre-processing, quality control, normalization and visualization of single-cell RNA-seq data in R. Bioinforma Oxf Engl. 2017;33(8):1179–1186. doi:10.1093/bioinformatics/btw777

42. Germain PL, Lun A, Meixide CG, Macnair W, Robinson MD. Doublet identification in single-cell sequencing data using *scDblFinder*. Published online May 16, 2022. doi:10.12688/f1000research.73600.2

43. Hao Y, Hao S, Andersen-Nissen E, et al. Integrated analysis of multimodal single-cell data. Cell. 2021;184(13):3573–3587.e29. doi:10.1016/j.cell.2021.04.048

44. Satija R, Farrell JA, Gennert D, Schier AF, Regev A. Spatial reconstruction of single-cell gene expression data. Nat Biotechnol. 2015;33(5):495–502. doi:10.1038/nbt.3192

45. Aran D, Looney AP, Liu L, et al. Reference-based analysis of lung single-cell sequencing reveals a transitional profibrotic macrophage. Nat Immunol. 2019;20(2):163–172. doi:10.1038/s41590-018-0276-y

46. Hevner RF. Layer-specific markers as probes for neuron type identity in human neocortex and malformations of cortical development. J Neuropathol Exp Neurol. 2007;66(2):101–109. doi:10.1097/nen.0b013e3180301c06

47. Lim L, Mi D, Llorca A, Marín O. Development and Functional Diversification of Cortical Interneurons. Neuron. 2018;100(2):294–313. doi:10.1016/j.neuron.2018.10.009

48. Sun W, Cornwell A, Li J, et al. SOX9 Is an Astrocyte-Specific Nuclear Marker in the Adult Brain Outside the Neurogenic Regions. J Neurosci. 2017;37(17):4493–4507. doi:10.1523/JNEUROSCI.3199-16.2017

49. Tremblay R, Lee S, Rudy B. GABAergic Interneurons in the Neocortex: From Cellular Properties to Circuits. Neuron. 2016;91(2):260–292. doi:10.1016/j.neuron.2016.06.033

50. Xin W, Mironova YA, Shen H, et al. Oligodendrocytes Support Neuronal Glutamatergic Transmission via Expression of Glutamine Synthetase. Cell Rep. 2019;27(8):2262–2271.e5. doi:10.1016/j.celrep.2019.04.094

51. Gautier O, Blum JA, Maksymetz J, et al. Human Motor Neurons Are Rare and Can Be Transcriptomically Divided into Known Subtypes. Neuroscience; 2023. doi:10.1101/2023.04.05.535689

52. Crowell HL, Soneson C, Germain PL, et al. muscat detects subpopulation-specific state transitions from multi-sample multi-condition single-cell transcriptomics data. Nat Commun. 2020;11(1):6077. doi:10.1038/s41467-020-19894-4

53. Risso D, Schwartz K, Sherlock G, Dudoit S. GC-content normalization for RNA-Seq data. BMC Bioinformatics. 2011;12:480. doi:10.1186/1471-2105-12-480

54. Robinson MD, McCarthy DJ, Smyth GK. edgeR: a Bioconductor package for differential expression analysis of digital gene expression data. Bioinforma Oxf Engl. 2010;26(1):139–140. doi:10.1093/bioinformatics/btp616

55. Bullard JH, Purdom E, Hansen KD, Dudoit S. Evaluation of statistical methods for normalization and differential expression in mRNA-Seq experiments. BMC Bioinformatics. 2010;11:94. doi:10.1186/1471-2105-11-94

56. Benjamini Y, Hochberg Y. Controlling the False Discovery Rate: A Practical and Powerful Approach to Multiple Testing. J R Stat Soc Ser B Methodol. 1995;57(1):289–300.

57. Conway JR, Lex A, Gehlenborg N. UpSetR: an R package for the visualization of intersecting sets and their properties. Bioinformatics. 2017;33(18):2938–2940. doi:10.1093/bioinformatics/btx364

58. Huang DW, Sherman BT, Lempicki RA. Systematic and integrative analysis of large gene lists using DAVID bioinformatics resources. Nat Protoc. 2009;4(1):44–57. doi:10.1038/nprot.2008.211

59. Sherman BT, Hao M, Qiu J, et al. DAVID: a web server for functional enrichment analysis and functional annotation of gene lists (2021 update). Nucleic Acids Res. 2022;50(W1):W216–W221. doi:10.1093/nar/gkac194

60. Ritchie ME, Phipson B, Wu D, et al. limma powers differential expression analyses for RNA-sequencing and microarray studies. Nucleic Acids Res. 2015;43(7):e47. doi:10.1093/nar/gkv007

61. Peixoto, Lucia H Stephanie, Risso, Davide. A Global Transcriptional Atlas of the Effect of Sleep Deprivation in the Mouse Frontal Cortex. Published online 2023. doi:10.5281/zenodo.10041833

