## Supplementary material for "A Global Transcriptional Atlas of the Effect of Sleep Deprivation in the Mouse Frontal Cortex": Inventory of Supplementary information

**Extended Data Figures:**

**Extended Data Figure 1. Table depicts the number of nuclei assigned to each cell-type for each mouse.** The first column depicts each cell-type (Astro, Car3, Endo, L2/3 IT CTX, L4/5 IT CTX, L5 IT CTX, L5 PT CTX, L5/6 NP CTX, L6 CT CTX, L6 IT CTX, L6b CTX, Lamp5, Micro-PVM, Oligo, Pvalb, SMC-Peri, Sncg, Sst, Sst Chodl, Vip, VLMC). Columns 2-7 depict each replicate, and the number of nuclei assigned to a given cell-type. Column 8 is the total number of nuclei in a given cell-type. HC, Home Cage Controls, SD, Sleep Deprived.

**Extended Data Figure 2. Validation of cell-type assignment pipeline using datasets from the BRAIN Initiative Cell Census atlas. A-C)** Uniform Manifold Approximation and Projection **(**UMAP) plot of the primary motor cortex (MOp) dataset (technique: 10X v3) from Allen Institute for Brain Science, colored according to **A)** dataset labels, **B)** Azimuth, **C)** SingleR labels. **D-F)** UMAP plot of the primary motor cortex (MOp) dataset (technique: 10X v2) from Allen Institute for Brain Science, colored according to **D)** dataset labels, **E)** Azimuth, **F)** SingleR labels.

**Extended Data Figure 3. Heatmap of cell-type specific marker expression in each cell group.** The average expression across all replicates (log-normalized counts) are shown for each marker in each cell-type. **A)** Heatmap with columns containing cell-types, and rows containing cell-type specific marker expression, unscaled **B)** Heatmap with columns containing cell-types, and rows containing cell-type specific marker expression scaled by rows to allow for row-to-row comparison.

**Extended Data Figure 4. The effect of sleep deprivation on non-neuronal cell-types and meta-analysis negative control genes. A)** Multidimensional scaling (MDS) plots depict the similarity of non-neuronal cell-types, including astrocytes, micro-PVM, oligodendrocytes and SMC-Peri. **B)** MDS plot verifies that previously published negative controls (Gerstner et al. 2016) are not affected by cell-type or condition. N=3 mice per condition. Home cage controls (HC) are shown in circles and sleep deprived (SD) animals are triangles.

**Extended Data Figure 5. Principal component analysis (PCA) plots for each cell-type following technical and biological normalization.** PCA plots following library size normalization with upper quartile and removal of unwanted variation via RUVs (k=2) from the RUVseq package. Home cage (HC) controls are shown in black circles, and sleep deprived (SD) animals are shown in red triangles. N=3 mice per condition. Only cell-types with more than 500 nuclei are included.

**Extended Data Figure 6. Volcano plots for each cell-type following differential gene expression (DGE) analysis with edgeR.** On the x-axis, fold-change is shown as logFC. On the y-axis, p-values are shown as -log10(p-value). Expressed genes are shown in gray. Significantly differentially expressed genes (DEGs) are shown in black, with the number of DEGs shown reported in the bottom left corner of each plot, FDR < 0.05. Positive controls from a previously published meta-analysis (Gerstner et al., 2016) are shown in red. Only cell-types with more than 500 nuclei are included. N=3 per condition.

**Extended Data Figure 7. Distribution of unadjusted p-values following pseudobulk differential gene expression (DGE) analysis with edgeR.** Only cell-types with more than 500 nuclei are included. N=3 per condition.

**Extended Data Figure 8. Simultaneous DGE and DTE analysis after SD. A-B)** PCA plots of counts following RUVs normalization (k=4) show sleep deprivation is the main source of variance in the data (first principal component, PC1) for both **A)** DGE, and **B)** DTE analyses with Swish. The percent variance is specified on each axis for the first and second source of variance (second principal component, PC2) in the data. Circles: HC, triangles: SD. **C-D)** Distribution of unadjusted p-values following **C)** DGE and **D)** DTE analyses with Swish. **E)** Bar plots depict the fold changes of upregulated (red) and downregulated (blue) eukaryotic initiation factor transcripts that are significantly differentially expressed, q-value < 0.05 and |log2FC| > 0.2. Transcripts in bold represent genes that are also significantly differentially expressed following DGE analysis, q-value < 0.05 and |log2FC| > 0.2. N=5 per condition. HC, Home Cage. SD, Sleep Deprivation.

**Extended Data Figure 9. Applying a log2FC filter optimizes the recovery of positive and negative control genes.** Total number and intersection of genes and transcripts recovered at log2FC thresholds, separated by upregulated and downregulated. For each threshold for gene level analysis, the rate of positive and negative controls recovered is reported. FC, fold change.

**Supplementary Tables:**

**Figure 2 Supplementary Table 1.** List of differentially expressed genes (SD compared to HC, FDR < 0.05) within cell-types with more than 500 nuclei, separated by tab. Color coded for upregulated (red) and downregulated (blue). SD, Sleep Deprived. HC, Home Cage Controls.

**Figure 2 Supplementary Table 2.** List of genes known to be affected by sleep deprivation. ‘Genes’ contains a full list of genes with references, ‘Subset’ contains a smaller list of ‘Genes.’

**Figure 3 Supplementary Table 1.** List of genes from the glutamatergic and GABAergic Venn diagram, separated by tab. LogFC for differentially expressed genes (FDR < 0.05) are shown. Color coded by upregulated (red) and downregulated (blue)**.**

**Figure 3 Supplementary Table 2.** Functional enrichment analysis output from DAVID for genes unique to glutamatergic or GABAergic neurons or shared between the two. Clustered terms are grouped together as they have a similarity threshold > 0.2, and enrichment score > 1.5. Color coded by upregulated (red) and downregulated (blue)

**Figure 4 Supplementary Table 1.** Lists of unique differentially expressed genes (SD compared to HC, FDR < 0.05) within cell-types with more than 500 nuclei, separated by tab. Color coded for upregulated (red) and downregulated (blue). SD, Sleep Deprived. HC, Home Cage Controls.

**Figure 4. Supplementary Table 2.** Functional enrichment analysis output from DAVID for genes unique to a cell-type. Clustered terms are grouped together as they have a similarity threshold > 0.2, and enrichment score > 1.5. Color coded by upregulated (red) and downregulated (blue).

**Figure 5 Supplementary Table 1.** List of differentially expressed genes and transcripts (SD compared to HC, q-value < 0.05), separated by tab and color coded for upregulated (red) and downregulated (blue). SD, Sleep Deprived. HC, Home Cage Controls.

**Figure 5 Supplementary Table 2.** List of positive control genes that are significantly differentially expressed for bulk gene level analysis (q-value < 0.05).

**Figure 6 Supplementary Table 1.** List of differentially used transcripts following sleep deprivation (SD compared to HC, q-value < 0.05). Color coded by increasing proportion (red) and decreasing proportion (blue). SD, Sleep Deprived. HC, Home Cage Controls.

**Figure 6 Supplementary Table 2.** Median normalized inferential replicate values, separated by tab for differential transcript expression and usage analyses. One value per biological replicate.

**Figure 7 Supplementary Table 1.** Functional enrichment analysis output from DAVID for the 1,575 genes that have significant differential transcript usage. Clustered terms are grouped together as they have a similarity threshold > 0.2, and enrichment score > 1.5.

Supplementary tables can be found at Zenodo with the following DOI: https://doi.org/10.5281/zenodo.10041833**.**
