## Extended Data Figures for "A Global Transcriptional Atlas of the Effect of Sleep Deprivation in the Mouse Frontal Cortex"

Extended Data Figure 1

| Cell-type | Mouse | | | | | | N. of nuclei |
|---|---|---|---|---|---|---|---|
| | 1 HC | 2 SD | 3 HC | 4 SD | 5 HC | 6 SD | |
| Astro | 142 | 151 | 87 | 122 | 260 | 285 | 1047 |
| Car3 | 17 | 44 | 37 | 58 | 77 | 19 | 252 |
| Endo | 21 | 181 | 33 | 1 | 9 | 114 | 359 |
| L2/3 IT CTX | 2844 | 2835 | 2496 | 2456 | 2330 | 2444 | 15405 |
| L4/5 IT CTX | 1717 | 981 | 1843 | 1520 | 1141 | 1140 | 8342 |
| L5 IT CTX | 673 | 707 | 791 | 641 | 381 | 571 | 3764 |
| L5 PT CTX | 557 | 574 | 638 | 825 | 637 | 565 | 3796 |
| L5/6 NP CTX | 63 | 88 | 68 | 102 | 165 | 153 | 639 |
| L6 CT CTX | 774 | 744 | 689 | 661 | 826 | 765 | 4459 |
| L6 IT CTX | 1185 | 1160 | 857 | 968 | 644 | 858 | 5672 |
| L6b CTX | 53 | 78 | 56 | 95 | 99 | 162 | 543 |
| Lamp5 | 20 | 31 | 42 | 54 | 126 | 51 | 324 |
| Micro-PVM | 77 | 60 | 41 | 62 | 79 | 260 | 579 |
| Oligo | 97 | 136 | 74 | 114 | 319 | 321 | 1061 |
| Pvalb | 368 | 320 | 367 | 457 | 428 | 457 | 2397 |
| SMC-Peri | 54 | 381 | 63 | 34 | 48 | 90 | 670 |
| Sncg | 78 | 72 | 57 | 67 | 51 | 71 | 396 |
| Sst | 239 | 200 | 265 | 326 | 529 | 378 | 1937 |
| Sst Chodl | 7 | 17 | 8 | 10 | 22 | 11 | 75 |
| Vip | 129 | 78 | 102 | 125 | 125 | 136 | 695 |
| VLMC | 14 | 67 | 24 | 22 | 82 | 30 | 239 |

Extended Data Figure 2

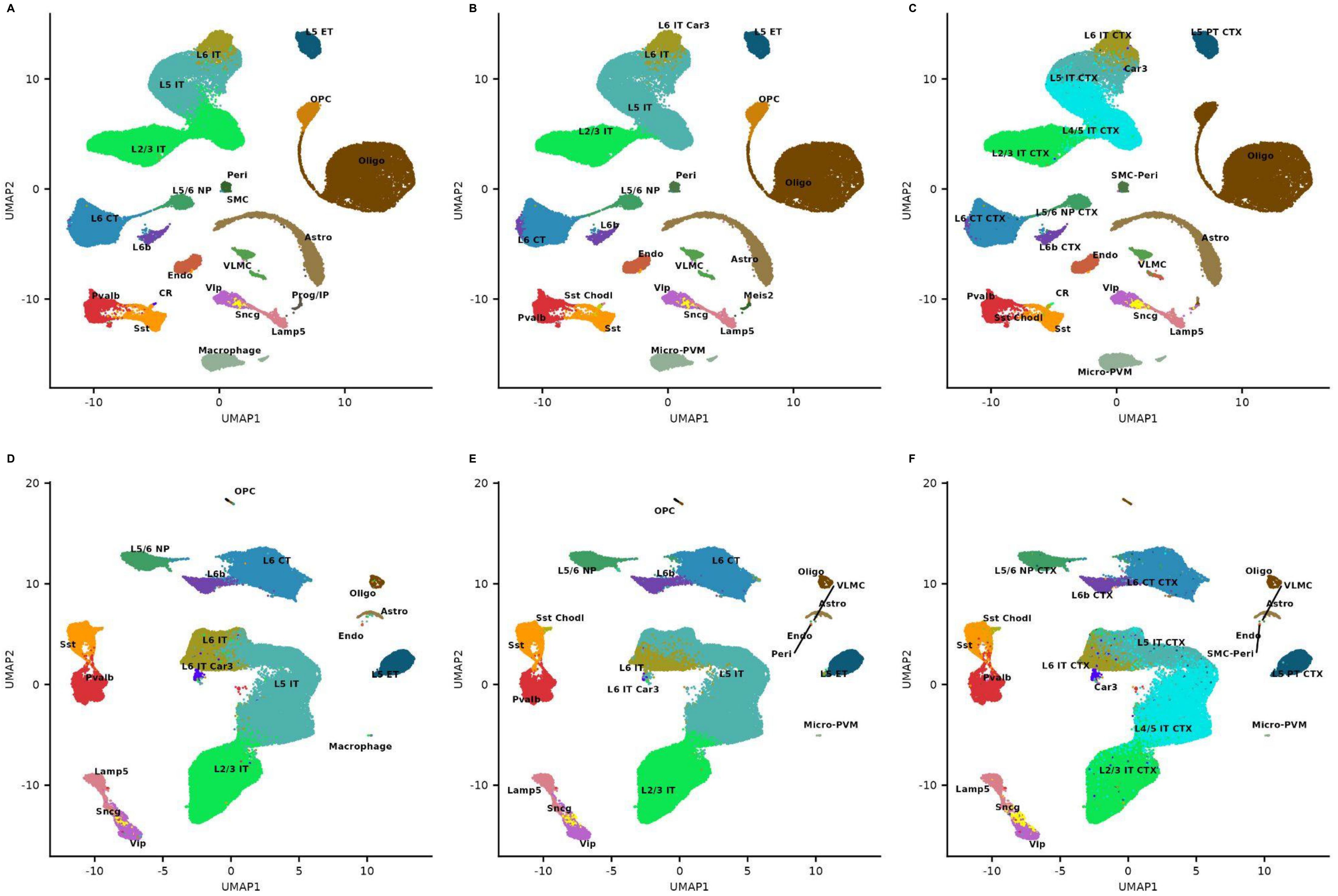

Extended Data Figure 3

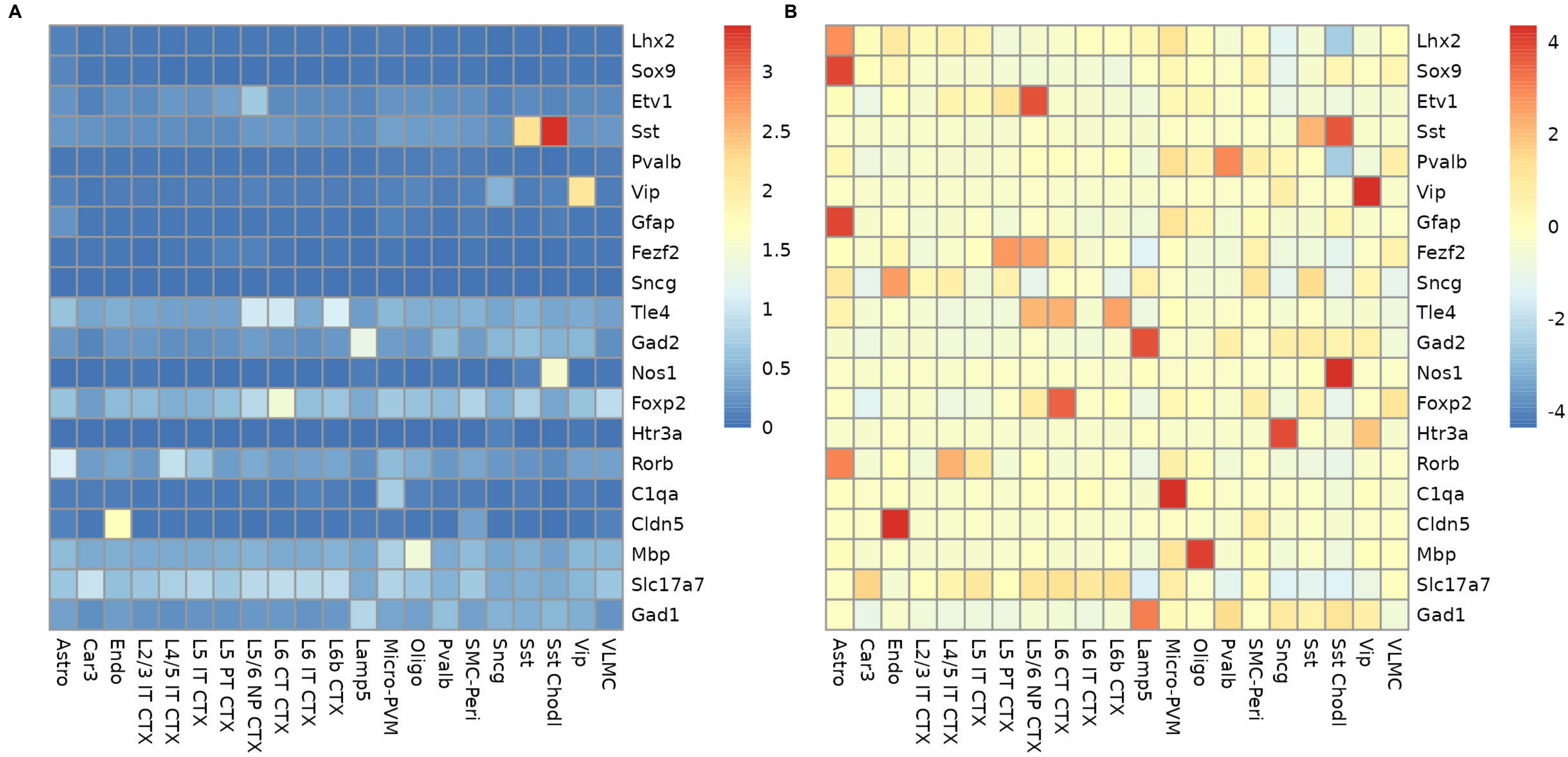

Extended Data Figure 4

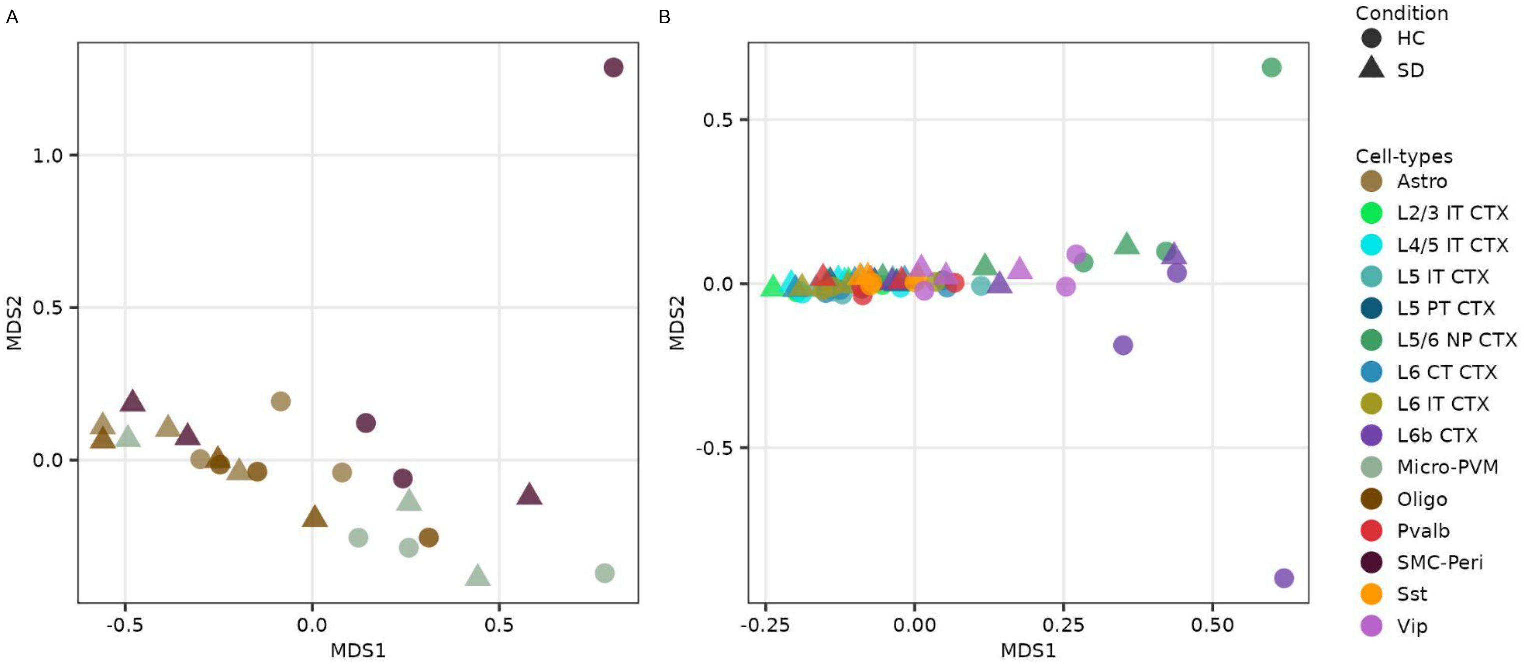

Extended Data Figure 5

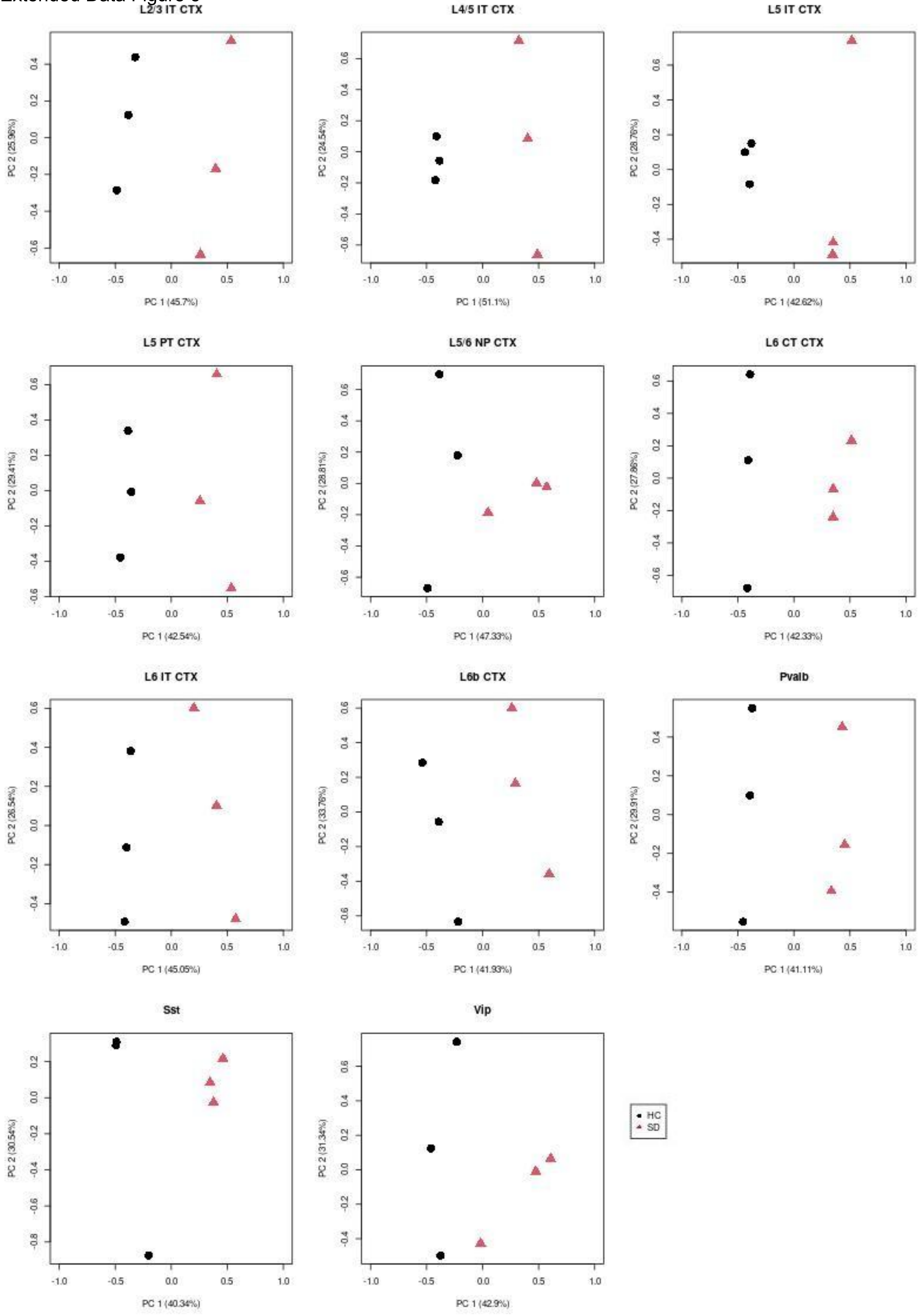

Extended Data Figure 6

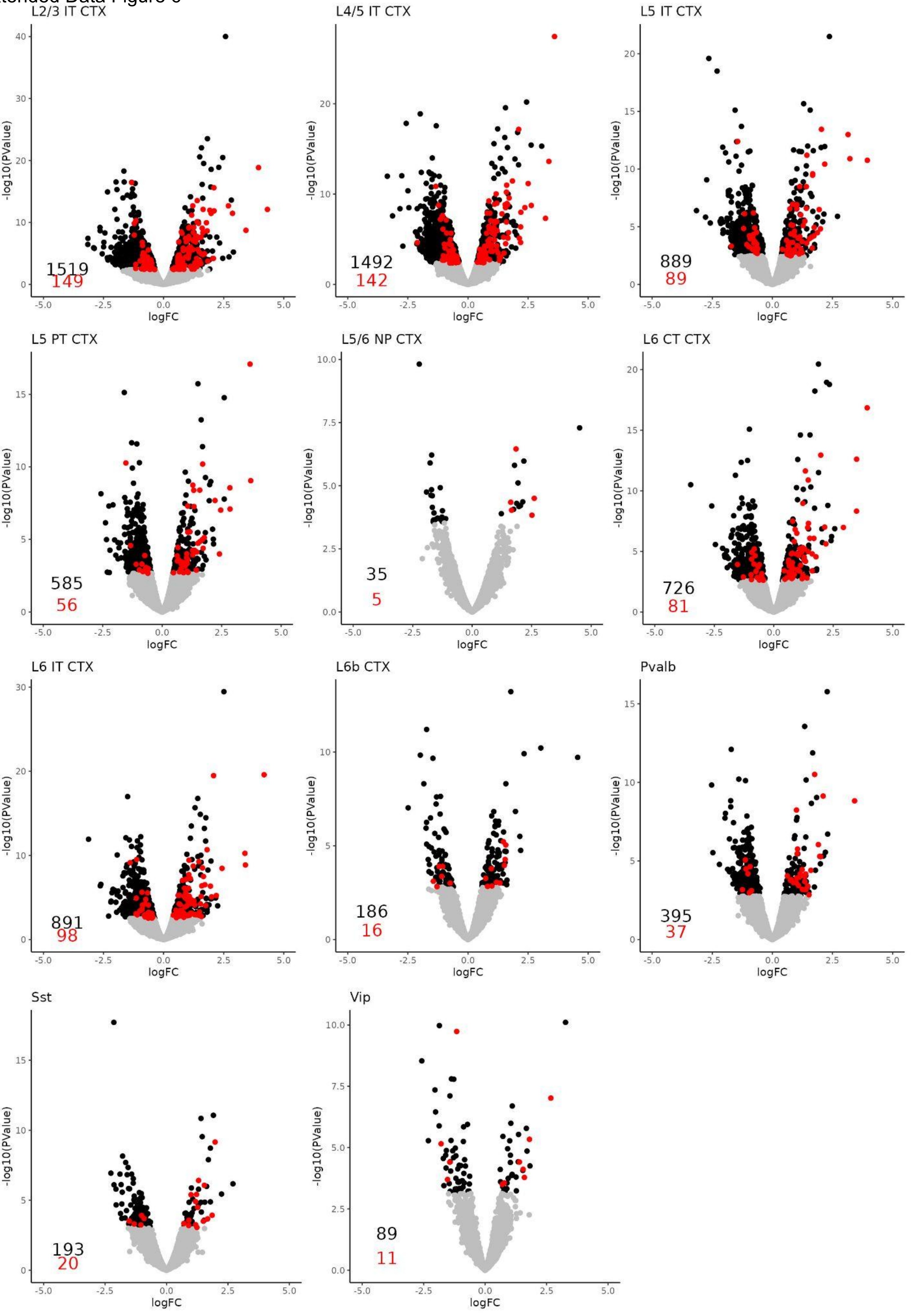

Extended Data Figure 7

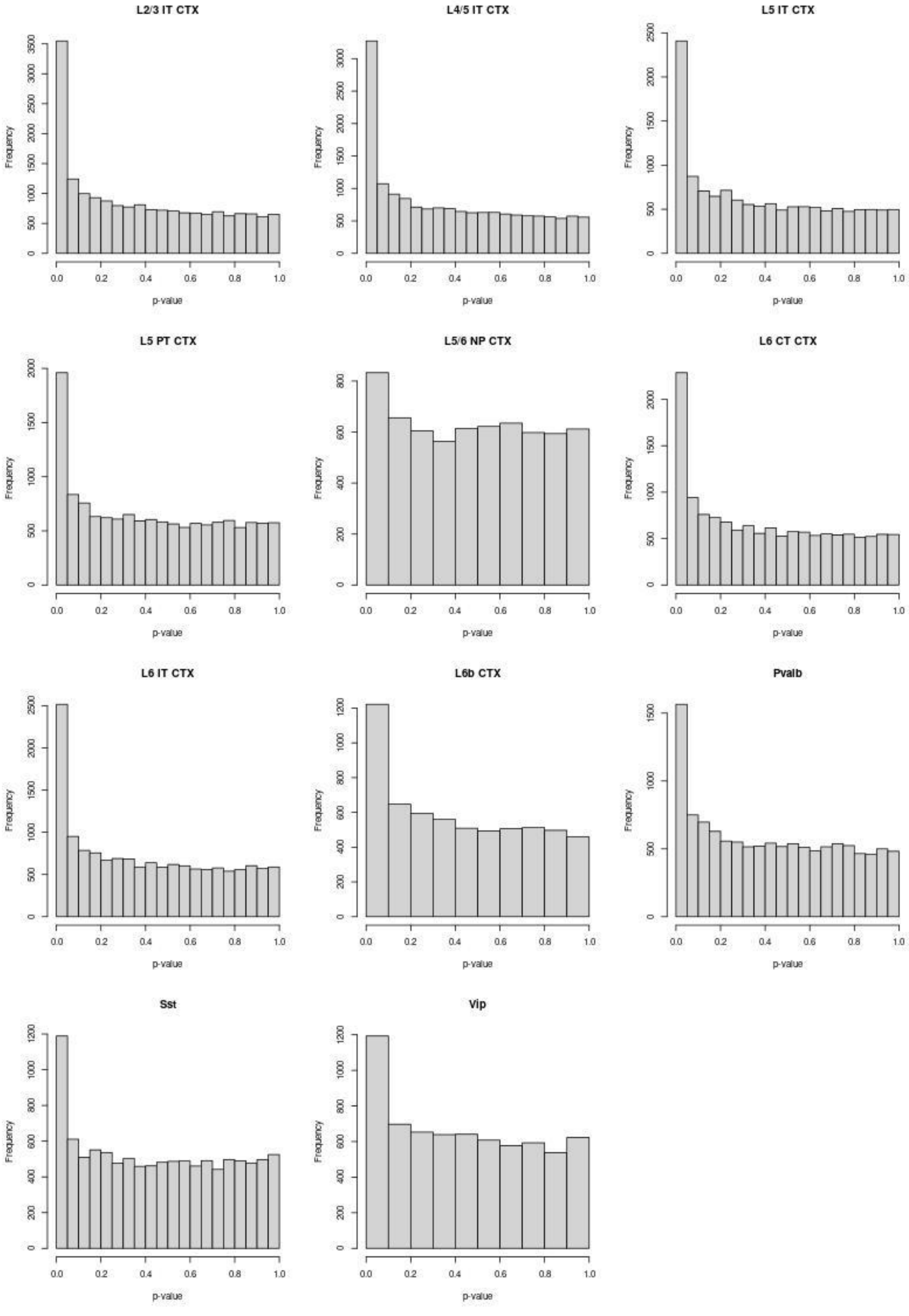

Extended Data Figure 8

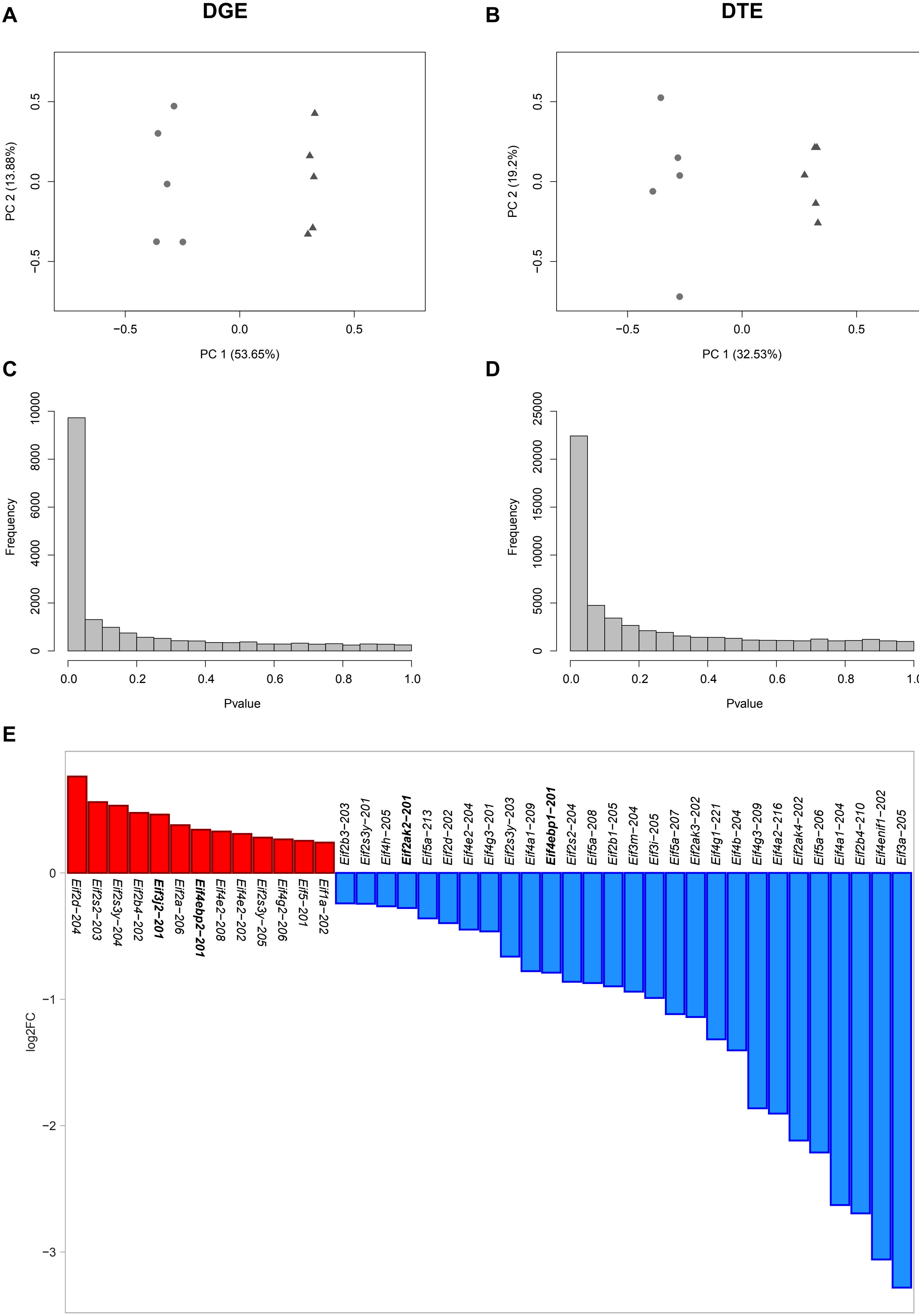

Extended Data Figure 9

| log2FC | | | | | | | | | | | | |
|---|---|---|---|---|---|---|---|---|---|---|---|---|
| **log2FC Cutoff (Absolute Value)** | **Number of Genes Detected by DGE Analysis (qvalue < 0.05)** | **Number of Genes Detected by DTE Analysis (qvalue < 0.05)** | **Up-regulated Intersections (qvalue < 0.05)** | | | **Down-regulated Intersections (qvalue < 0.05)** | | | **Positive Controls Recovered (Gene)** | | **Negative Controls Recovered (Gene)** | |
| | | | **DTE, Not DGE** | **DGE, Not DTE** | **Intersection of DGE and DTE** | **DTE, Not DGE** | **DGE, Not DTE** | **Intersection of DGE and DTE** | **#** | **% (Relative to Expressed Matrix)** | **#** | **% (Relative to Expressed Matrix)** |
| 0 | 8505 | 10439 | 1381 | 872 | 3269 | 3117 | 528 | 3836 | 558 | 83.2 | 1309 | 43.3 |
| 0.1 | 7506 | 9687 | 1272 | 691 | 2899 | 3001 | 447 | 3469 | 535 | 79.7 | 1105 | 36.5 |
| **0.2** | **4863** | **7341** | **920** | **381** | **1819** | **2724** | **290** | **2373** | **411** | **61.3** | **592** | **19.6** |
| 0.3 | 3027 | 5482 | 638 | 219 | 1061 | 2510 | 205 | 1542 | 294 | 43.8 | 318 | 10.5 |
| 0.4 | 1984 | 4270 | 412 | 137 | 675 | 2303 | 145 | 1027 | 214 | 31.9 | 188 | 6.2 |
| 0.5 | 1308 | 3367 | 272 | 86 | 445 | 2060 | 105 | 672 | 157 | 23.4 | 124 | 4.1 |
